# Tropical rainforest *versus* savannah: Biome-specific landscape structure mediates adaptive strategies and resilience to environmental change

**DOI:** 10.64898/2026.09.21.753388

**Authors:** Katie Gates, Jonathan Sandoval-Castillo, Chris J. Brauer, Peter J. Unmack, Martin Laporte, Louis Bernatchez, Luciano B. Beheregaray

**Affiliations:** Molecular Ecology Laboratory, College of Science and Engineering, Flinders University, Adelaide, SA 5042, Australia; Institute for Applied Ecology, University of Canberra, ACT 2601, Australia; Institut de Biologie Intégrative et des Systèmes (IBIS), Université Laval, 1030 avenue de la Médecine, Québec, G1V 0A6, Canada; Ministère de l’Environnement, de la Lutte contre les changements climatiques, de la Faune et des Parcs, 880 chemin Sainte-Foy, Québec, G1S4X4, Canada

**Keywords:** landscape genomics, climate adaptation, freshwater fish, tropical biodiversity, genotype-environment association, climate change

## Abstract

As ecosystems face rapid environmental change, understanding the interplay between landscape structure, genetic diversity, and adaptive potential is critical for predicting resilience. We investigated how biome-specific connectivity and environmental heterogeneity mediate adaptive strategies and resilience in the tropical rainbowfish (*Melanotaenia splendida splendida*) across contrasting rainforest and savannah ecosystems in tropical Australia. Using an integrative framework combining genomic, morphological, and environmental analyses, we inferred that hydroclimatic variation has been a key driver of genetic and phenotypic divergence. Rainforest populations exhibited strong local adaptation to fragmented, stable environments, while savannah populations showed less genomic differentiation and morphological divergence, patterns consistent with weaker local adaptation. We hypothesise that greater reliance on phenotypic plasticity may facilitate adaptive responses in savannah populations, reflecting their exposure to more connected and variable environments. Variance partitioning highlighted the contrasting roles of neutral and adaptive processes across biomes, with implications for managing resilience in freshwater systems under climate change. Our findings underscore the trade-offs between connectivity, local adaptation, and plasticity, providing a model for understanding adaptation and resilience across diverse ecosystems.

## Introduction

Understanding relationships between adaptive diversity and environment will be a prerequisite for anticipating and managing ecological responses to climate change in coming decades. The importance of climate as a selective force is well established (Franks and Hoffmann 2012, Anderson and Song 2020), and where migratory opportunities are limited, such as for freshwater organisms, patterns of standing adaptive diversity are likely to be an important determinant of local resilience (Sgrò et al. 2011). Such adaptive patterns are influenced by the selective environment in which a species evolved (Holderegger et al. 2006, Whitlock 2014), and may therefore vary widely in accordance with local or regional conditions (Moritz et al. 2012), in addition to demographic and life history traits (Clarke 1979). Consequently, it is expected that broader patterns of resilience are also likely to vary geographically, influenced by factors such as local and regional climatic variability, and the strength of ecological gradients (Deutsch et al. 2008, Tewksbury et al. 2008). It has been suggested that tropical regions may be more vulnerable to changing climates because of organisms’ narrow thermal niches (Huey et al. 2009, Sunday et al. 2011), limited thermal tolerance and plastic abilities (Holzmann et al. 2026). This may be particularly apparent for ectotherms such as freshwater fishes due to their limited internal thermoregulatory capacities, and to projected changes in water flow, water temperature extremes and riverine community abundance and structure (Rohr and Palmer 2013, Barbarossa et al. 2021, Brown et al. 2024). However, very little is known about the extent that ecological adaptation contributes to tropical diversity or about the adaptive relevance of climatic variation across different tropical habitats.

Transition zones such as the interface between rainforest and savannah are particularly promising arenas for the study of environmental adaptation and resilience (Smith et al. 1997, Ostridge et al. 2025). Rainforest and savannah are the most dominant biomes in the terrestrial tropics, varying not only climatically, but in the structural and functional complexity of their biotic communities (Murphy and Bowman 2012, Bond et al. 2021). Although rainforest and savannah often occur adjacently, most species distributions are non-overlapping, reflecting conflicting habitat requirements (Fensham 1995, Azihou et al. 2013). While both are highly biodiverse, rainforests are typically taxonomically richer (Ter Steege et al. 2000, Kier et al. 2005) and include a greater proportion of obligate associations (Fensham 1995, Ibanez et al. 2013). A history of climatic fluctuations and frequent fire activity has contributed to greater temporal variability of savannah communities (Staver et al. 2011b, Kutt et al. 2012, Vasconcellos et al. 2019). In contrast, many rainforest regions have experienced long-term climatic continuity, maintaining a proliferation of ancient lineages. This stability is one factor thought to have promoted the accumulation of tropical diversity and specialisation (Gaston and Blackburn 1996, Kooyman et al. 2013). However, landscape and environmental heterogeneity must also be considered if we are to adequately explain diversification in tropical bioregions (Moritz and McDonald 2005, Dagallier et al. 2020, Furness et al. 2021).

To formulate hypotheses about likely adaptive influences across rainforest and savannah, we can consider the bioclimatic interactions that are consistently associated with bioregion boundaries. Savannah communities are typically more dominant where annual rainfall is less than ∼1,000 mm, and rainforest where more than ∼2,000 mm (Hirota et al. 2011, Staver et al. 2011a). Meanwhile, rainfall, fire activity, and substrate may be subsequently influenced by forest density, producing feedback loops which help to sustain distributions (Hirota et al. 2011, Oliveras and Malhi 2016, Wu et al. 2016). These factors have broader implications for organisms’ exposure to annual, seasonal, and diurnal climatic extremes, whereby savannah organisms are subjected to more variable and greater intensities of most climatic variables than in the rainforest. Notably, regional bioclimatic dynamics also appear to be influenced by topography, with rainforest biotas more often occurring in rugged and complex terrain (Murphy and Bowman 2012, Ondei et al. 2017). This could in some cases contribute to greater microhabitat structure and less landscape connectivity in rainforests (Svenning 1999), affecting the spatial scale over which both neutral and adaptive divergences may occur (Nosil et al. 2019). It is therefore possible that rainforest and savannah organisms may differ not only in response to regional climatic influences, but in the extent of locally specific adaptation within biomes, both with likely flow-on effects to resilience in changing conditions.

Disentangling environmental influences on adaptive and non-adaptive variation in wild populations is greatly assisted by genomic datasets, which are expected to encompass loci varying both neutrally and in response to selective pressures (Holderegger et al. 2006, Schwartz et al. 2010). Moreover, if ecotypic adaptations are a result of heritable evolutionary changes, then relevant associations with environment are likely to be reflected by both genomic and physiological divergence (Santure and Garant 2018). Landscape genomics approaches are increasingly seeking to identify overlap between genotype-environmental associations and divergence in fitness-related traits (Balkenhol et al. 2017, Bernatchez et al. 2024). However, a limited number of studies have so far explicitly tested genotype-phenotype-environment associations in natural populations (Vangestel et al. 2018, Smith et al. 2020). This may not only provide a more holistic approach for assessing relevant environmental influences, but also improve inferences about candidate genes underlying environmental and climatic resilience (Carvalho et al. 2021). Additionally, large discrepancies in morphological and genetic patterns may highlight a reliance on plasticity for physiological changes, while strong overlaps can further support evolutionary responses to selection (Merilä and Hendry 2014).

Australian rainbowfishes (*Melanotaenia*) are a valuable system to study climatic adaptation in freshwater ecosystems (McCairns et al. 2016, Gates et al. 2017, Brauer et al. 2018, Sandoval-Castillo et al. 2020, Smith et al. 2020, Attard et al. 2022, Brauer et al. 2023). Previous rainbowfish studies have shown that body shape divergence has evolved repeatedly in response to hydroclimatic selection (McGuigan et al. 2003, McGuigan et al. 2005, Sandoval-Castillo et al. 2020) and provided evidence for adaptive (genetic) plasticity due to ecotype-specific directional selection (Sandoval-Castillo et al. 2020). Furthermore, this group includes examples of relatively recent divergence of species and subspecies across ecological transitions (McGuigan et al. 2000, Hurwood and Hughes 2001, Unmack et al. 2013). In a tropical context, where ancient lineages proliferate, this creates an opportunity for comparative evolutionary studies assessing ongoing mechanisms of divergence (Moritz et al. 2000). Like many Australian rainbowfishes, tropical-endemic *Melanotania splendida splendida* exhibits extensive phenotypic variation across their range, which includes both rainforest and savannah ecoregions. Morphological, meristic and colour variations have been observed between populations, drainages (i.e. river catchments), and even contrasting habitats within drainages (Pusey et al. 2004). This has led to suggestions of substantial within-species genetic diversity and/or a highly variable and plastic phenotype (Pusey et al. 2004). Within the species’ rainforest distribution, evidence was provided for hydroclimate-associated genomic and morphological variation (Gates et al. 2023). Moreover, these environmental influences could account for a greater proportion of biological variation than measures of neutral divergence. This suggests local adaptation has been highly relevant to the intraspecies diversity, with implications for additional adaptation required to withstand climatic changes (Gates et al. 2023). The addition of savannah representatives is therefore desirable for assessing broader patterns of trait divergence in tropical landscapes, which is expected to be influenced by both local and regional adaptation. The species’ rainforest distribution is relatively rugged and topographically complex, being to a large extent determined by the presence of the highlands of the Great Dividing Range (Nott 2005). Rainforest drainage networks are ancient, densely packed, and mostly perennial (Nott 2005, Pearson 2005, Pearson et al. 2015). In contrast, streams across the lowland drainage systems of Cape York’s savannah regions are often ephemeral, but connect at greater spatial scales due to branching between major tributaries during high volume monsoonal runoff (Howley et al. 2013). This comparative scenario provides the opportunity to assess influences of both the hydroclimate and terrain structure on adaptive variation and resilience.

To this end, we used a landscape genomics approach to test environmental associations with genotype, phenotype, and genotype-phenotype interactions in *M. s. splendida*, both between and within tropical biomes. Given the striking climatic and ecological differences between ecoregions, we hypothesised that the greatest intraspecies divergence may also occur across the rainforest-savannah interface. We also predicted that environmental associations could be better at explaining biological variation than neutral factors, especially for body shape variation which was inferred to have important functional relevance. Within ecoregions, we also interrogated the effects of terrain connectivity on spatial patterns of adaptation, asking whether gene flow may act to strengthen or homogenise local signals of genetic and morphological adaptation. Addressing these questions is important for understanding the dynamics of evolution in tropical freshwaters and may inform prioritisation of management strategies under rapid climatic change.

## Methods

### Field sampling

We collected wild *Melanotaenia splendida splendida* (eastern rainbowfish) from seventeen sites in tropical north-eastern Australia in March 2017. Localities included nine rainforest creek sites across five drainages, and eight savannah creek sites across one drainage (Figure 1; Table 1). A total of 510 individuals were captured using seine nets and euthanised on the day of capture via overdose of anaesthetic sedative (AQUI-S^®^: 175mg/L, 20 minutes) at mobile fieldwork stations, followed immediately by digital photographing for morphometrics (final photographic dataset of 366 individuals, avg. 22, min. 13 per sampling site; Table 1). Fin clips from all collected samples were preserved in 99% ethanol and stored at -80°C. Of these, 381 high quality samples were chosen for the final DNA dataset (avg. 22, min. 15 per site; Table 1). For 302 individuals (avg. 18, min. 11 per site), final genomic and morphometric datasets overlapped, enabling direct contrasts in later association tests among genotype, phenotype, and environment.

**Figure 1.**
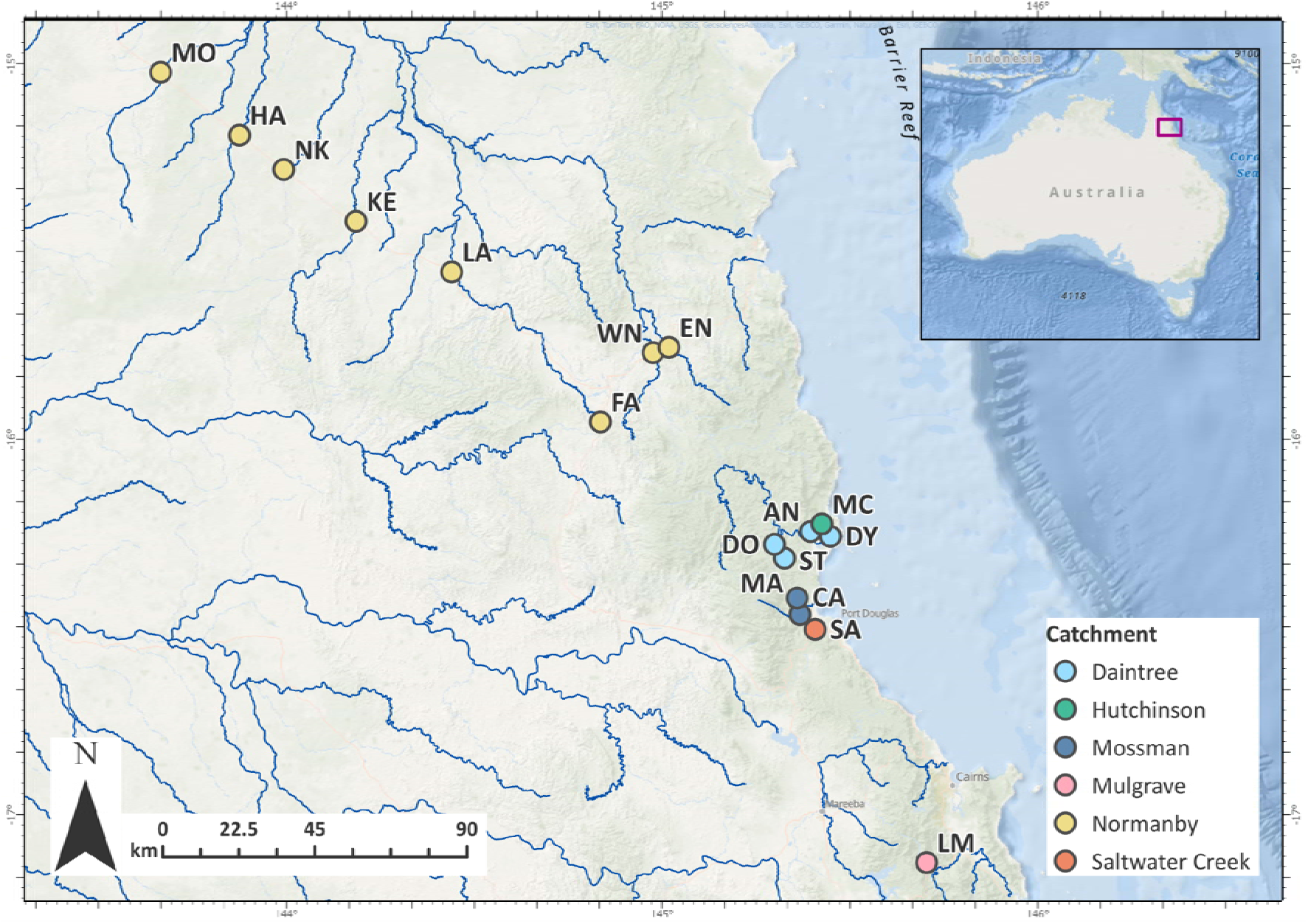
Sampling location map of *Melanotaenia splendida splendida* collected from eight savannah (shown in yellow) and nine rainforest locations (other colours) in north-eastern Australia. Point colours correspond to river drainages of origin; all savannah locations were sampled in the Normanby River drainage. Navy lines represent major river channels within the region. Inset: extent indicator of main map relative to the Australian continent.

**Table 1.** Localities and sample sizes (*n*) of *Melanotaenia splendida splendida* collected from rainforest and savannah biomes of tropical north-eastern Australia for genomic and morphometric data.

| Location | Site Code | River Drainage | Latitude | Longitude | Ecotype | Total collected n | Final n (DNA) | Final n (Morpho) | Final n (GxPxE) |
| --- | --- | --- | --- | --- | --- | --- | --- | --- | --- |
| Little Mulgrave Creek | LM | Mulgrave | -17.13 | 145.7 | Rainforest | 30 | 23 | 20 | 17 |
| Cassowary Creek | CA | Mossman | -16.51 | 145.41 | Rainforest | 30 | 23 | 30 | 23 |
| Marrs Creek | MA | Mossman | -16.47 | 145.36 | Rainforest | 24 | 20 | 19 | 15 |
| Saltwater Creek | SA | Saltwater Creek | -16.42 | 145.36 | Rainforest | 30 | 24 | 21 | 19 |
| Stewart Creek | ST | Daintree | -16.32 | 145.32 | Rainforest | 30 | 25 | 22 | 20 |
| Douglas Creek | DO | Daintree | -16.28 | 145.3 | Rainforest | 30 | 24 | 29 | 21 |
| Doyle Creek | DY | Daintree | -16.26 | 145.45 | Rainforest | 30 | 24 | 23 | 22 |
| Forest Creek | AN | Daintree | -16.25 | 145.39 | Rainforest | 31 | 22 | 21 | 18 |
| McClean Creek | MC | Hutchinson | -16.23 | 145.42 | Rainforest | 32 | 25 | 22 | 22 |
| Famechon Creek | FA | Normanby | -15.95 | 144.83 | Savannah | 30 | 25 | 19 | 17 |
| West Normanby River | WN | Normanby | -15.77 | 144.97 | Savannah | 30 | 21 | 20 | 18 |
| East Normanby River | EN | Normanby | -15.76 | 145.02 | Savannah | 30 | 19 | 13 | 11 |
| Laura River | LA | Normanby | -15.56 | 144.44 | Savannah | 33 | 23 | 22 | 11 |
| Kennedy River | KE | Normanby | -15.42 | 144.18 | Savannah | 30 | 21 | 20 | 16 |
| North Kennedy River | NK | Normanby | -15.28 | 143.99 | Savannah | 30 | 15 | 20 | 15 |
| Hann River | HA | Normanby | -15.19 | 143.87 | Savannah | 30 | 24 | 23 | 18 |
| Morehead River | MO | Normanby | -15.02 | 143.66 | Savannah | 30 | 23 | 22 | 19 |

### Genomic data collection

DNA was extracted from fin clips by salting-out, using a protocol modified from Sunnucks and Hales (1996). We then assessed DNA quality, quantity, and integrity using NanoDrop (Thermo Scientific), Qubit (Life Technologies), and gel electrophoresis (agarose, 2%) respectively. We produced double-digest restriction site-associated DNA (ddRAD) libraries in-house following Peterson et al. (2012) with modifications according to Sandoval Castillo et al. (2018) for 420 individuals (including replicates and those later removed during filtering). We assigned samples randomly over five sequencing lanes (∼six replicates per lane), of which four were sequenced by the South Australian Health and Medical Research Institute Genomics Facility (Illumina HiSeq25000; single-ended), and one by Novogene Hong Kong (Illumina HiSeq4000; paired-ended).

Raw sequences were demultiplexed and trimmed of adaptors and leading/trailing low quality bases (Phred < 20) using TRIMMOMATIC 0.39 within the DDOCENT 2.2.19 pipeline (Puritz et al. 2014). Poorly sequenced individuals (< 700,000 reads) were removed from the dataset. Sequences were mapped to a rainbowfish reference genome (*M. duboulayi*; NCBI GenBank accession number JAPDEC000000000) using the GATK 3.7 pipeline (Van der Auwera and D O’Connor 2020). The SNP variants were called from the mapped reads using BCFTOOLS 1.9 (Li 2011), and filtered using VCFTOOLS 0.1.15 (Danecek et al. 2011) to remove poorly sequenced reads, non-biologically informative artefacts (*sensu* O’Leary et al. (2018)), complex variants, and sites with high likelihood of linkage. For the full filtered dataset, we used BAYESCAN 2.1 (Foll and Gaggiotti 2008) to assess locus-specific conformity to neutral expectations based on allele frequency distributions across populations (with population membership first inferred by preliminary FASTSTRUCTURE 1.0 (Raj et al. 2014). We used BAYESCAN default settings and a false discovery rate of < 0.05, producing a putatively neutral dataset of 14,479 SNPs for subsequent analyses of neutral genetic diversity and population structure.

### Genomic diversity and inferences of population structure

Locality-specific neutral genomic diversity was assessed using ARLEQUIN 3.5 (Excoffier and Lischer 2010) to determine mean expected heterozygosity (*H_e_*), mean nucleotide diversity (π), and proportion of polymorphic loci (*PP*). Pairwise *F*_ST,_ site-specific *F*_ST_, and global *F*-statistics were calculated in R (RC Team 2019) using HIERFSTAT 0.04-22 (Goudet 2005). The latter were calculated for all individuals (‘between-systems’), as well as independently within each ecoregion (‘savannah-specific’ and ‘rainforest-specific). Additionally, we produced a scaled covariance matrix of population allele frequencies (Ω) using BAYPASS 2.2 (Gautier 2015) core model, based on all SNPs rather than the neutral subset, which is implicitly estimated. This hierarchical Bayesian model provides an informative basis for demographic inference by accounting for structure resulting from shared history. The method follows from the BayEnv model proposed by (Coop et al. 2010, Günther and Coop 2013), but with several extensions to improve accuracy by estimation of prior distributions. These were plotted in R using BAYPASS’s included utility functions. We further interrogated population structure using a FASTSTRUCTURE clustering approach, also repeated both between and within ecoregions (see Gates et al. (2023) for full description of methods).

### Characterising environmental variation

We used the same six attributes from the Australian Hydrological Geospatial Fabric (Geoscience Australia 2011; Stein (2011) from previous rainforest-specific analyses (Gates et al. 2023). Given that covariation among variables differed depending on region and on spatial scale, we were able to include two additional variables in between-systems analyses, and were required to use one fewer in savannah-specific analyses. For between-systems analyses, we included stream segment aspect (ASPECT), river disturbance index (RDI), mean summer runoff (RUNSUMMERMEAN), mean winter runoff (RUNWINTERMEAN), mean annual rainfall (STRANNRAIN), mean annual temperature (STRANNTEMP), total length of upstream segments calculated for the segment pour-point (STRDENSITY), and stream segment slope (VALLEYSLOPE) (Supplementary Table 1). Rainforest-specific analyses excluded RUNWINTERMEAN and VALLEYSLOPE, while savannah-specific analyses excluded ASPECT, STRANNRAIN and VALLEYSLOPE. Environmental variables were used as a basis for genotype-environment associations (GEA), phenotype-environment associations (PEA) and genotype-phenotype-environment (GxPxE) associations, as described below.

### Geometric morphometric characterisation and analyses

We used TPSDIG2 2.31(Rohlf 2017) to position eighteen landmarks on the field-collected digital photographs. Landmarks (Supplementary Figure 1) were chosen to maximise homology, repeatability, and putative ecological relevance. In MORPHOJ 1.07a (Klingenberg 2011), digitised TPS files were compiled and subjected to Procrustes superimposition, screened for outliers representing landmarking errors, and used to produce Procrustes covariance matrices for ‘between-systems’, ‘rainforest-specific’ and ‘savannah-specific’ subsets. To characterise major features of shape variation, PCAs were performed on resulting matrices. Allometric regressions, pooled within populations identified in neutral genetic analyses, were used to determine a positive association between size (log centroid) and shape (Procrustes coordinates). Regression residuals were therefore used to test for relationships between body shape and locality. To for test ecoregional differences in the ‘between-systems’ dataset, we used a discriminant function analysis classifying by rainforest and savannah origin, with 1000 permutation rounds. Additionally, we used a canonical variate analysis (CVA) to determine variation among sampling sites, scaling for relative within-group variation. We again used 1000 permutations to test significance.

### Detecting selection between and within ecoregions

To test for environmental adaptation and divergence both between and within ecoregions, we used a combination of GEA, PEA, and GxPxE approaches. In all instances, we used partial redundancy analyses (pRDAs) in the R package VEGAN 2.5-6 (Oksanen et al. 2019). For GEAs only, we also incorporated a Bayesian hierarchical model (BAYPASS 2.2 (Gautier 2015)), which is tailored to genetic analysis and is well suited to study systems with hierarchical population structure (Gautier 2015).

For the GEAs, we first ran a global RDA using the full dataset (14,478 SNPs) for all individuals as a multivariate response matrix, and the eight ‘between-systems’ environmental variables described above as an explanatory matrix, which was first centred and scaled. Then, using only the environmental explanatory variables found to be associated in the global model (*p* = <0.1), and again using the full set of SNP genotypes as a response matrix, we repeated the analysis using three partial RDAs to control for putative neutral influences. In each, we included a different neutral (or neutral proxy) covariable matrix: 1) significant principal components (PCs) of scaled population allelic covariance (Ω), 2) significant PCs of pairwise *F*_ST_, and 3) scaled river distances. Further details of input file creation are in Gates et al. (2023), except for the river distances covariable, for which pairwise distances among connected sites were calculated in ARCMAP 10.3 (ESRI 2011), and distances between unconnected sites (i.e. different drainages) were imputed with distances an order of magnitude higher than the average. Starting from the global RDA, these steps were then repeated for ‘savannah-specific’ and ‘rainforest-specific’ analyses, including individuals and covariables specific to the region. Finally, we used the alternative method of BAYPASS 2.2 to produce ‘savannah-specific’, ‘rainforest-specific’ and ‘between-systems’ GEA analyses, using the auxiliary covariate model with default settings and the same sets of scaled environmental explanatory variables as for the pRDAs. Here, we accounted for assumed population demographic structure via the scaled covariance matrix of population allele frequencies (Ω) resulting from the core model.

For the PEAs, we adapted the same pRDA approach, with the same environmental explanatory datasets, to test for signals of selection in the observed morphological variation. Here, the response matrix comprised PCs of individual Procrustes distances determined significant by Broken-Stick modelling, again controlled for putatively neutral genetic structure (allelic covariance Ω; pairwise *F*_ST_; river distances), plus the additional covariable of body size (log centroid size). Inputs for the body shape response variable and size covariable were created in _R_, using functions developed by Claude (2008). Although sexual dimorphism may produce an additional confounding effect on body shape variation, we found that equal sex ratios were present between sampling regions (11:14 m:f, Chi-Square *p* value = 0.987) and we therefore did not include sex as a covariable. As with GEAs, PEA analyses were repeated for ‘between-systems’, ‘rainforest-specific’ and ‘savannah-specific’ datasets.

### Genotype-phenotype-environment analysis

We used a GxPxE approach to test whether environmentally associated genetic variation could be attributed to morphological adaptation between and within ecoregions. Using R, we ran a global RDA using significant PCs of individual Procrustes distances as explanatory variables, and putative adaptive alleles (candidates combined from genotype-environment RDAs controlling for Ω and BAYPASS 2.2 auxiliary covariate model, described above) as the multivariate responses. The analysis was then repeated as a partial RDA using individual body size (log centroid) as a covariable. This enabled isolation of only the genotype-phenotype associations best explained by environmental selection, and with the potential to underlie heritable body shape variation.

## Results

### Sequencing, bioinformatics, genetic diversity and population structure

Filtering of genome-mapped sequencing reads produced a putatively unlinked dataset of 14,540 SNPs, of which 14,478 were considered neutral for the purposes of population genomic analyses. The full and neutral datasets comprised 381 individuals across nine sampling sites. We found moderately high neutral genomic diversity (Table 2) with expected heterozygosity (*H*_E_) among sites ranging from 0.278 to 0.321 (mean = 0.289), and proportion of polymorphic loci (*PP*) ranging from 0.252 to 0.395 (mean = 0.349). Site-specific averages were similar between rainforest and savannah systems, with *H*_E_ slightly higher in the rainforest (rainforest mean = 0.293; savannah mean = 0.284), and *PP* slightly higher in the savannah (rainforest mean = 0.329; savannah mean = 0.372). However, ranges of variation for all diversity measures were greater among rainforest sites.

**Table 2.**
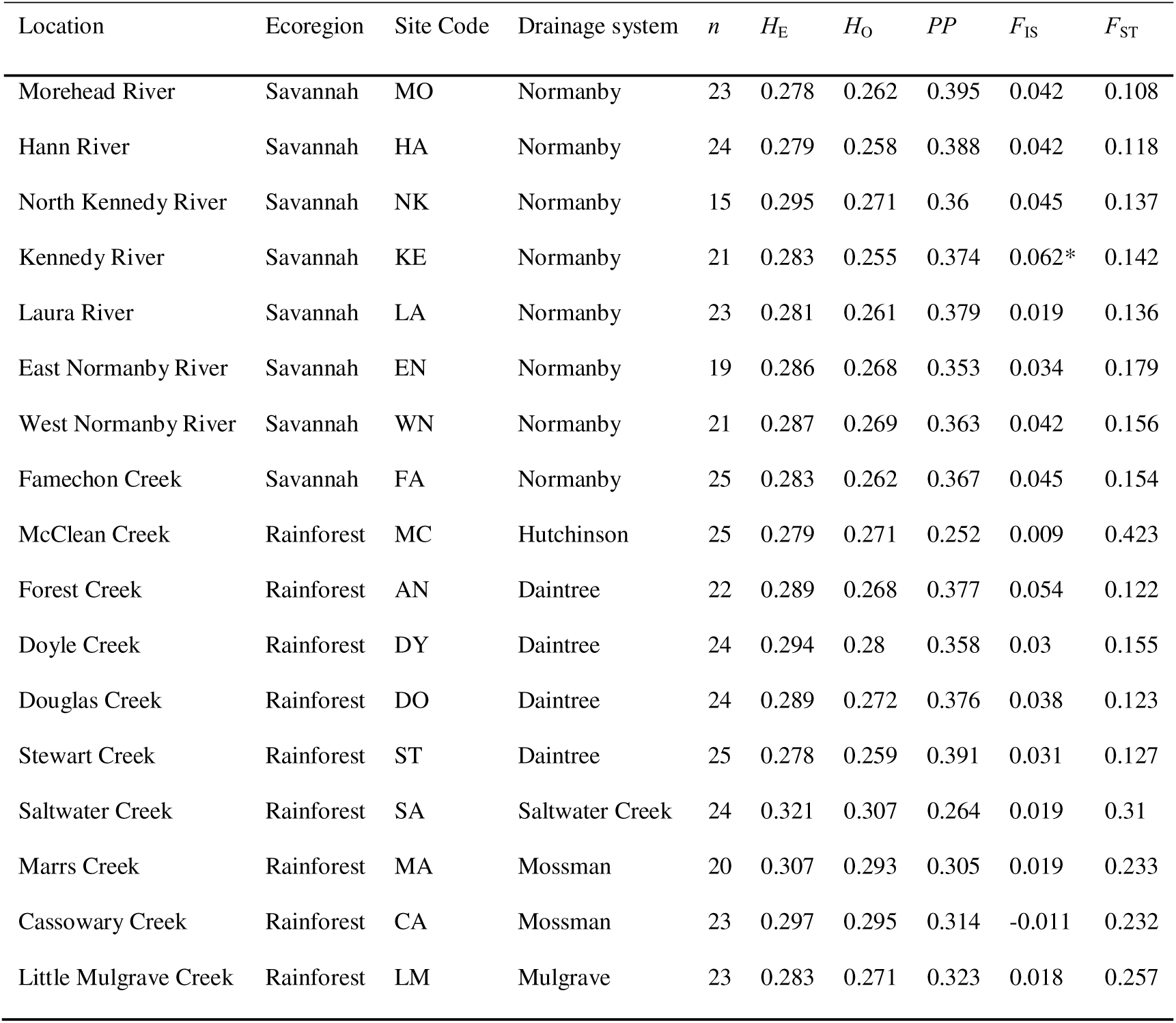
Genetic diversity measures for the eastern rainbowfish *Melanotaenia splendida splendida* at nine rainforest and eight savannah localities, based on 14,478 putatively neutral loci (n = sample size for final DNA dataset; *H*_E_ = expected heterozygosity; *H*_O_ = observed heterozygosity; *PP* = proportion of polymorphic loci; *F*_IS_ = site-specific inbreeding coefficient (values with *p* = < 0.05 indicated by *); *F*_ST_ = site-specific *F*_ST_).

Global *F*_ST_ values were higher in the rainforest system (0.148) than in the savannah (0.025) (Global *F*_ST_ between-systems was 0.173). Site-specific (Table 2) and pairwise *F*_ST_ values (Supplementary Table 2) indicated that much of this divergence could be attributed to inter-drainage rather than intra-drainage differences. This was also reflected by the stronger correlations in allelic covariance within river drainages, detected by BAYPASS (Supplementary Figure 2). Global *F*_IS_ was slightly higher in the savannah (0.0741) than the rainforest (0.0490). Clustering analyses also indicated that the main neutral genetic differentiations occurred among river drainages. In between-systems FASTSTRUCTURE analysis (Figure 2a), individuals were grouped by drainage with exception of those from Saltwater; these were grouped together with those from the neighbouring Mossman drainage, under a best *K* of five. However, in ecoregion-specific analyses (Figure 2b), rainforest alone was found to have an optimal *K* of five. Very little admixture was visible between Saltwater and Mossman, indicating hierarchical substructure may have obscured differentiation in the combined-systems analysis. An optimal *K* of one was found in the savannah-specific analysis of the single Normanby drainage, however Figure 2c displays *K* = 2, to demonstrate regional substructure. Locality-based clustering using singular value decompositions of Ω (Supplementary Figure 3) also showed strong separation between Saltwater and Mossman, even in combined-systems analysis. Therefore, contemporary evolutionary processes are likely occurring relatively independently among six drainage-associated subunits, which we will refer to as populations. These contrasting population structure patterns are consistent with lower connectivity and more independent evolutionary dynamics among rainforest drainages, and with higher connectivity and gene flow across the lowland savannah drainage network.

**Figure 2.**
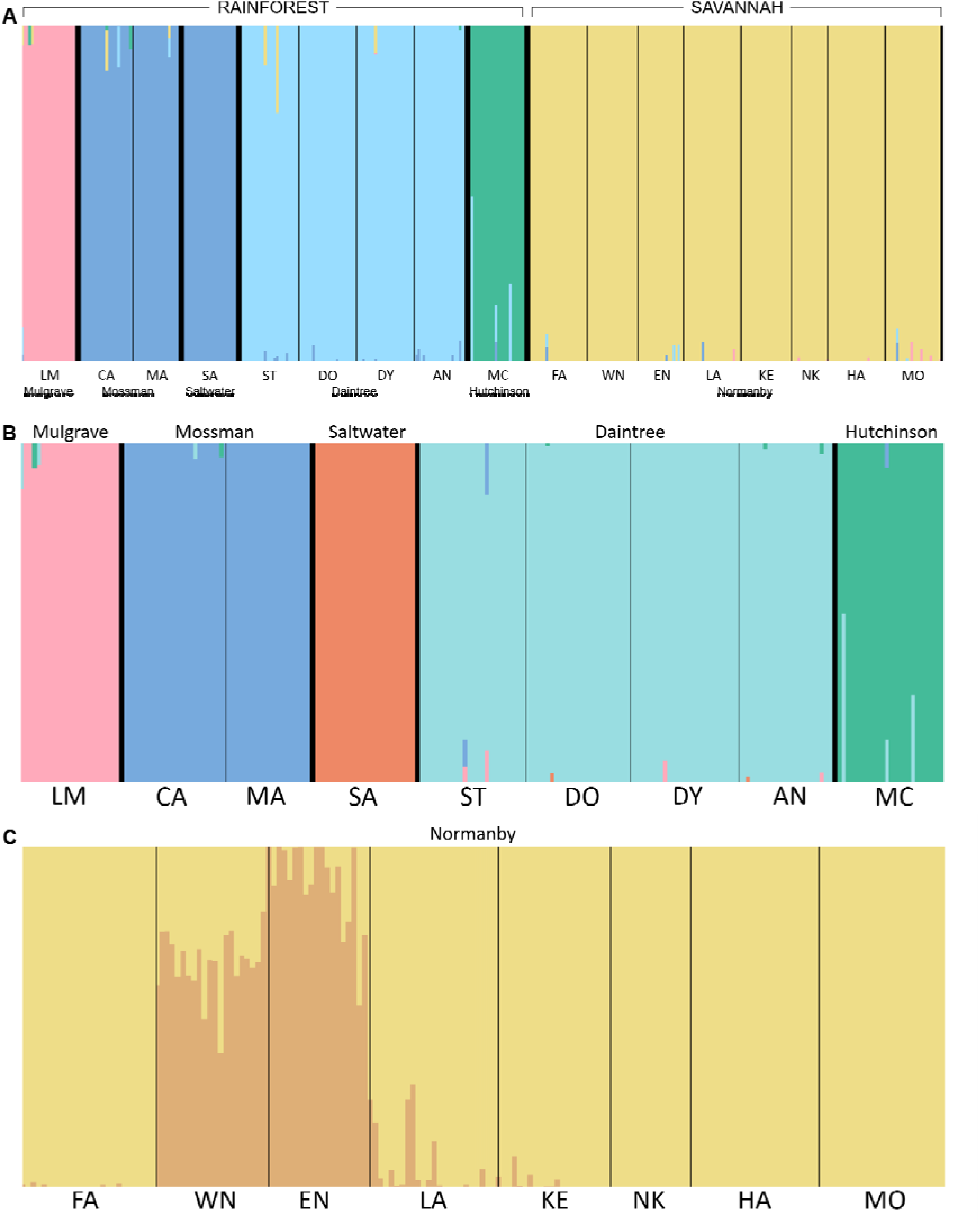
Cluster plots based on FASTSTRUCTURE analysis of 14,478 putatively neutral SNPs, where colours represent inferred ancestral populations of individuals based on A) all sampled individuals, showing optimal *K* = 5; B) only rainforest individuals, showing optimal *K* = 5; C) only savannah individuals, showing *K* = 2 to inform about regional substructure within the drainage system (actual inferred optimal *K* = 1). Large type refers to drainage systems, which are separated by thicker black lines. Small type refers to sampling localities, separated by thinner black lines. Locality abbreviations follow Table 1.

### Morphological divergence between ecoregions and sampling localities

The PCA of body shape of all *M. s. splendida* individuals produced four significant PCs under broken stick modelling (Supplementary Figure 4). Major aspects of variation included body depth (PC1), dorsal height and head orientation (PC2), length of caudal fork and caudal peduncle (PC3), width and position of fin bases (PCs 3 & 4), and size of eye (PC4). Site-specific CVAs found significant differences in mean body shape among most localities after controlling for size (*p* < 0.05), but with substantial overlap among individuals from different localities (Supplementary Figure 5; Supplementary Table 3). However, strong separation between rainforest and savannah was evident on the first axis. Congruently, discriminant function analysis between rainforest and savannah individuals found that body shape could reliably classify individuals to ecoregion of origin in 96.5% of cases (94.3% in cross-validation; *p* = <0.0001; Supplementary Figure 6; Supplementary Table 4). Rainforest fish were larger on average than savannah fish (mean centroid size 10.20 cm (SD = 2.88 cm) in rainforest versus 6.93 cm (SD = 2.06 cm) in savannah), however they were also dorsoventrally narrower. Even after controlling for allometric differences, body depth was the most notable component of shape divergence between ecoregions (Figure 3).

**Figure 3.**
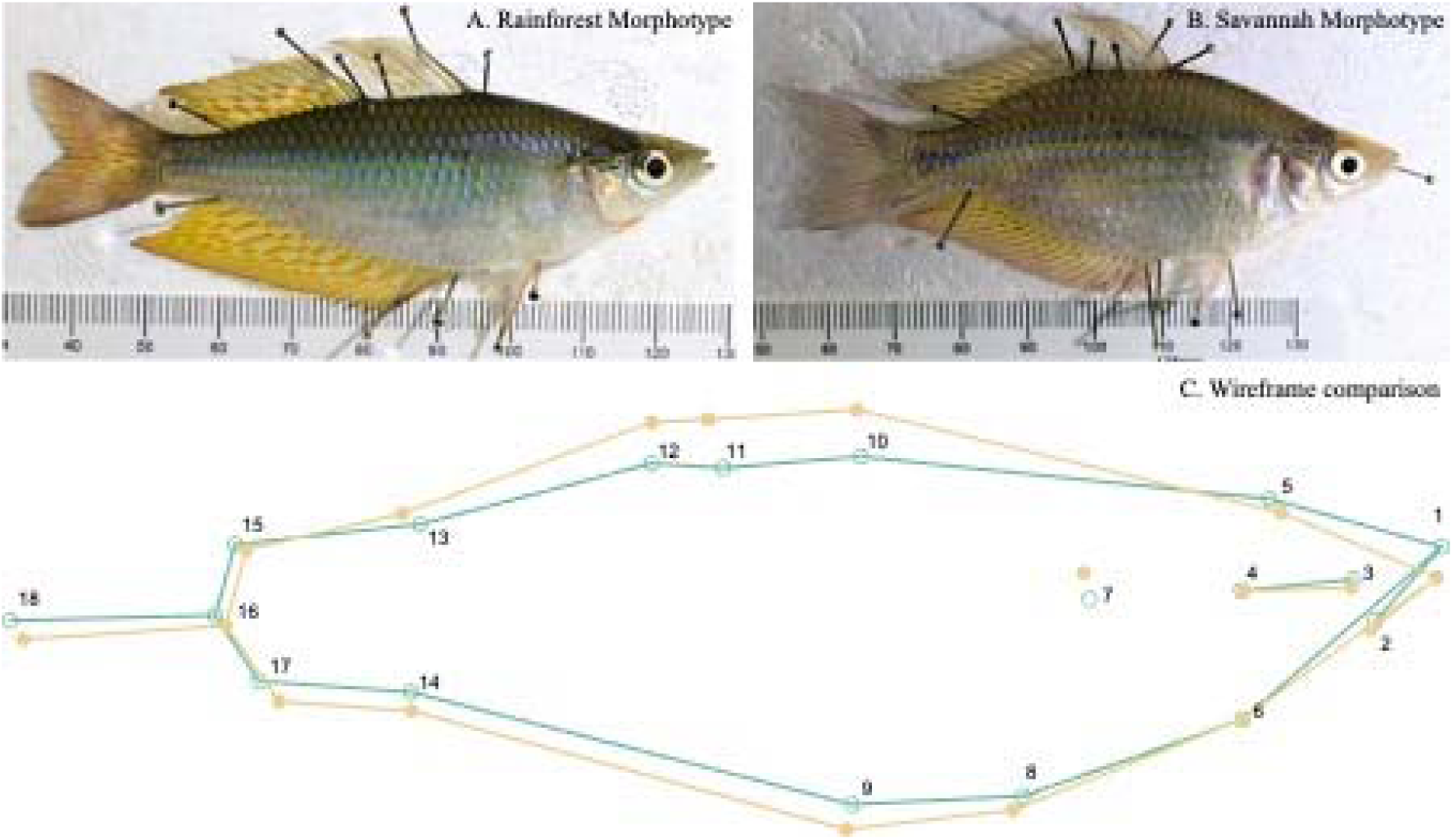
Morphological differences between *Melanotaenia splendida splendida* of rainforest and savannah origin. Photographs show examples of similarly sized males collected from rainforest (top left; Saltwater Creek (SA13); centroid size 12.55cm) and savannah (top right; Morehead Creek (MO01); centroid size 12.24 cm) in March, 2017. Wireframe diagram shows group mean shape change between rainforest and savannah individuals, based on discriminant function analysis of size regression residuals (18 landmarks, n = 367). Scale factor = 2. Green = rainforest. Yellow = savannah.

### Genomic and morphological associations with environment

The pRDA analyses found strong genomic and morphological associations with environment (Supplementary Figure 7; Supplementary Table 5), which consistently explained more of the observed biological variation than neutral or spatial factors (including *F*_ST_ distance, allelic covariance & river distance; Figure 4). On average, environment best explained 3.3 times more genomic variation and 10.5 times more morphological variation than other covariables (Supplementary Table 6).

**Figure 4.**
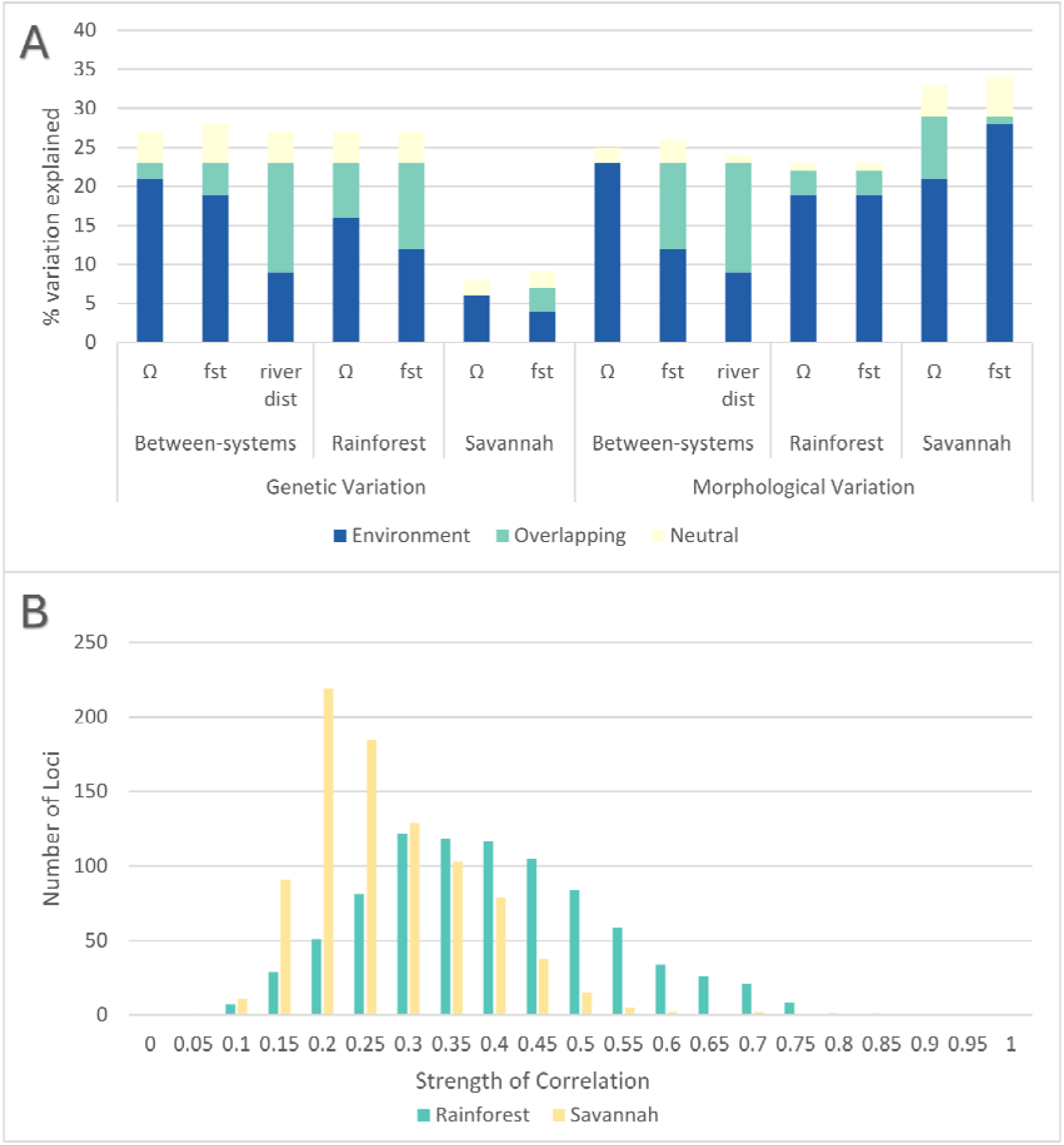
Relative contributions of environmental and neutral components to genomic and morphological variation in *Melanotaenia splendida splendida*. A) Variance partitioning of pRDA response variables, including genomic and morphological variation, showing the percentage of variation explained by environmental predictors, neutral covariables, or their shared components. Neutral covariables comprised allelic covariance (Ω), pairwise *F*_ST_ distances, and river distances. B) Frequency distribution of locus-specific correlations with environment for candidate adaptive SNPs identified in rainforest and savannah populations after controlling for putative neutral variation using allelic covariance (Ω). Ecoregion-specific pRDAs were based on the full dataset of 14,540 SNPs and detected 864 candidate SNPs in rainforest populations and 880 in savannah populations.

For between-systems analyses, the most important environmental explanatory variables for both genomic and morphological variation were average annual rainfall (STRANNRAIN) and summer mean runoff (RUNSUMMERMEAN). These were also the best explanatory variables identified by the alternative GEA approach of BAYPASS (auxiliary covariate model; Supplementary Figure 8). In rainforest-specific analyses, STRANNRAIN and average annual temperature (STRANNTEMP) were the best explanatory variables for both genomic and morphological variation. However, within the savannah, winter mean runoff (RUNWINTERMEAN) and STRANNTEMP best explained genetic variation, in contrast to morphological variation, which was best explained by RUNSUMMERMEAN and stream density (STRDENSITY). Both pRDAs and BAYPASS approaches produced suites of candidate genes for environmental adaptation, totalling 1,284 in between-systems analyses (1,119 pRDA, 233 BAYPASS, 68 shared), 1,004 in rainforest-specific analyses (864 pRDA, 176 BAYPASS, 36 shared), and 987 in savannah-specific analyses (880 pRDA, 145 BAYPASS, 38 shared). Although slightly more candidates were detected by pRDAs in the savannah compared to the rainforest, we found that locus-specific selective signals were weaker, with an average correlation of 0.249 in the savannah compared to 0.371 in the rainforest (Figure 7).

The pRDA associations among genotype, phenotype, and environment (Figure 5g-i) revealed significant relationships both between and within ecoregions (*p* = <0.001), even after controlling for size. Between-systems, 7.2% of environment-associated genetic variation could be explained by the first three PCs of body shape variation, revealing 212 SNPs as candidates for climate-adaptive morphological variation. Within the rainforest, 61 candidates were identified in association with the first four body shape PCs, while within the savannah, 72 candidates were identified in association with PCs 1, 2 and 4.

## Discussion

We compared strength and variation of adaptive diversity in native rainbowfish ecotypes to understand evolutionary divergence and provide a foundation for further considerations of climatic resilience across contrasting tropical freshwater biomes. Our results support a central role for hydroclimate in driving intraspecies divergence across rainforest and savannah, while differences in drainage structure and inferred gene flow appear to modulate the spatial scale of adaptation and resilience. Across systems, we found strong evidence for biome-specific hydroclimatic adaptation, and likely functional relevance of major morphological differences. Within systems, we also found high explanatory power of hydroclimatic variables in shaping local variation. However, regional genomic signals indicated a homogenising effect of gene flow on adaptive variation in the relatively well-connected savannah, fuelling the hypothesis of compensatory reliance on phenotypic plasticity in fluctuating environments. Our results indicate that differences in habitat variability and complexity can substantially affect evolutionary trajectories of tropical organisms, which we discuss in the context of resilience to a rapidly changing climate.

### Environmental and neutral contributions to intraspecies divergence

Despite their proximity, the major biomes of north-eastern Australia differ markedly in climate (Ash 1988). Wet tropical rainforests are relatively stable and aseasonal, whereas savannahs experience strong thermal and hydrological variability (Ash 1988, Bowman et al. 2010). Within-biome gradients are subtler but still substantial (Supplementary Figure 1).

Consistent with our first hypothesis, the strongest biological differences in *M. s. splendida* occurred between ecoregions, with genomic and morphological associations exceeding neutral expectations. Significant associations within both biomes further indicate adaptation at local and regional scales. Supporting previous work within the rainforest (Gates et al. 2023), environmental factors explained genomic and morphological variation better than neutral structure in both rainforest and savannah-specific analyses, and across combined systems. This pattern held across multiple pRDA models using different neutral covariables (Ω, FST, river distance). For morphology, environmental effects were especially strong, explaining approximately ten times more variation than neutral factors, compared to approximately three times for genomic data. This apparent resistance of phenotypes to neutral influences, especially relative to genomic patterns, is consistent with the hypothesis that body shape has been more constrained by functional requirements than has genomic variation. As expected, a substantial proportion of unexplained variation likely reflects within-site differences and stochastic processes not captured by the models.

Genomic variation was more strongly associated with neutral structure than was morphological variation, likely because many genomic loci are neutral or nearly neutral, even under divergent selection (Ohta 2002, Luikart et al. 2018). By contrast, integrative traits such as morphology may be more tightly linked to fitness (Ho et al. 2017, Zhang 2018). Complex traits like body shape interact across multiple biological levels, not just with the surrounding environment but with lower level components such as cells, tissues and organs (Zhang 2018). This potentially strengthens selection and inhibits the effect of drift on morphological adaptation. Consistent with significant G×P×E associations, morphological variation in rainbowfishes appears at least partly heritable, with plasticity likely further contributing (e.g., Kelly 2019).

### Putative selective influences and outcomes

Although studies of natural selection in tropical wild populations remain limited (Siepielski et al. 2017), it appears that hydrological, thermal, and vegetation gradients act as key selective drivers in terrestrial fauna (Ntie et al. 2017, Zhen et al. 2017, Miller et al. 2020, Morgan et al. 2020, Bennett et al. 2021). Here, and perhaps unsurprisingly for tropical aquatic obligates (Cooke et al. 2012, Cooke et al. 2014, Hay et al. 2022), we found that species-wide adaptive signals were most strongly linked to hydrology. Across both between-system environmental association analyses (pRDA and BAYPASS), average annual rainfall best explained genomic and morphological variation, followed by summer runoff. These variables aligned with pRDA axes and with rainforest–savannah divergence in both datasets. Within systems, hydroclimatic variation remained important, but annual variables (rainfall, temperature) best explained rainforest variation, whereas seasonal runoff dominated in savannah sites. This likely reflects stronger monsoonal seasonality in savannah environments (Ma et al. 2013). Our findings align with meta-analytical evidence that precipitation is a major driver of selection globally (Siepielski et al. 2017). Agreement between RDA and BAYPASS further supports these associations (Forester et al. 2018).

Morphologically, rainforest and savannah fish differed most in body and caudal peduncle depth. Rainforest individuals were larger but more streamlined, with dorsoventral flattening and narrower peduncles, whereas savannah fish were deeper-bodied with dorsal humps. These traits are functionally linked to swimming performance (Gatz 1979). Slender forms generally favour sustained swimming and high-flow environments, while deep bodies enhance manoeuvrability, burst swimming, and performance in low or variable flows (Gatz 1979, Scarnecchia 1988, Leavy and Bonner 2009, Langerhans and Reznick 2010, Alexandre et al. 2014). These patterns suggest that greater flow variability in savannah habitats may select for deeper bodies that enhance unsteady locomotion, particularly during low-flow periods, whereas rainforest conditions favour streamlined forms for efficient sustained swimming in consistently high flows (Langerhans 2008). However, evidence for body-depth adaptation in rainbowfishes is mixed across species, sexes, and environments (McGuigan et al. 2003, Lostrom et al. 2015, Kelley et al. 2017), so while hydrological adaptation is plausible, the underlying mechanisms remain unresolved. Even so, the combined genomic, morphological, and environmental associations indicate that hydroclimatic variation is an important axis of ecological divergence in this system. More broadly, these findings support the view that tropical diversity reflects both historical biogeographic processes and contemporary ecological divergence (Moritz et al. 2000, Moritz and McDonald 2005, Ricklefs 2005, Beheregaray et al. 2015). Given the strong hydroclimatic associations observed here, climate change may have substantial consequences for the distribution of adaptive diversity, particularly where local adaptation requires evolutionary turnover to maintain fitness under changing conditions (Fitzpatrick and Keller 2015, Bay et al. 2017).

### Adaptive dynamics under contrasting terrain structure

Climatic variability differs across Australia’s north-eastern tropical biomes, but rainforest and savannah regions are also separated by contrasting geomorphology, allowing assessment of how connectivity shapes neutral and adaptive divergence. *Melanotaenia s. splendida* from the five sampled rainforest drainages formed five distinct populations, with mild intra-drainage structure typical of moderately dispersing fishes (sensu Brauer et al. 2018). Although drainage networks may have retained similar terrain for tens of millions of years (Nott 2005), connectivity among these systems is likely more recent, for example via coastal floodplain exposure during glacial periods (Pusey and Kennard 1996, Cook and Hughes 2010), consistent with moderate pairwise *F*_ST_ values. Despite stronger clustering among neighbouring systems, there was little evidence of recent admixture, as might be expected under substantial gene flow during cyclonic rainfall (e.g. Pearson 2005) or via human translocations. In contrast, all savannah individuals were assigned to a single population, consistent with higher gene flow across connected lowland rivers. As in the rainforest, some intra-drainage substructure was detected, likely reflecting low winter flows, isolation by distance across large catchments, and multiple headwaters.

Connectivity differences between rainforest and savannah regions likely have broad impacts on freshwater biota. In fishes, drainage connectivity strongly influences species distributions, richness, and genetic diversity (Pusey and Kennard 1996, Unmack 2001, Wong et al. 2004, Carvajal-Quintero et al. 2019), and gene flow across heterogeneous landscapes can shape adaptive evolution (Garant et al. 2007, Nosil 2012, Tigano and Friesen 2016). Here, signals of local adaptation were consistently weaker in the savannah across both ecoregion-specific and combined GEA analyses, despite similar environmental variation. Savannah pRDA models showed lower explained variance, weaker clustering of adaptive variation, and fewer locus–environment associations than rainforest models. While this could reflect selection on many small-effect loci (Pritchard and Di Rienzo 2010), the total number of candidate associations was also lower. Consistently, the alternative GEA method BAYPASS identified fewer adaptive candidates in the savannah, and G×P×E pRDAs explained less variance and showed weaker clustering than in the rainforest.

Together, these results support our hypothesis that greater drainage connectivity in the savannah promotes homogenisation of adaptive variation, leading to more region-wide rather than locally specific genomic adaptation. In contrast, natural fragmentation in rainforest systems appears to enable more independent, locally driven adaptation among demes. This pattern aligns with expectations (sensu Slatkin 1987) that gene flow can dampen local divergence. Although gene flow can also facilitate adaptation (Tigano and Friesen 2016, Nosil et al. 2019), homogenisation here suggests it may be too high in the savannah for selection to counteract the swamping of adaptive alleles (sensu Storfer and Sih 1998). Alternatively, large population sizes of M. s. splendida in both regions (Pusey et al. 2004) may maintain sufficient standing variation for adaptation even in isolated rainforest drainages (sensu Jensen and Bachtrog 2011).

Differences in adaptive dynamics are likely to shape evolutionary responses to climate change, depending on spatial and temporal variability. In well-connected systems like the savannah, local adaptation may be constrained unless conditions push the system past a selective “tipping point,” after which gene flow can facilitate rapid, widespread change (Nosil et al. 2019). In contrast, poorly connected systems may show steadier, locally driven responses, but rely on independent emergence of adaptive variation within each deme (Tigano and Friesen 2016, Nosil et al. 2019), which may be limiting under rapid change (Brauer and Beheregaray 2020). These patterns suggest trade-offs between local specialisation and system-wide resilience in rainforest versus savannah populations. Given strong evidence for climatic divergence in *M. s. splendida* and other tropical species (Ntie et al. 2017, Zhen et al. 2017, Miller et al. 2020, Morgan et al. 2020, Bennett et al. 2021, Brauer et al. 2023), such trade-offs are likely important for long-term persistence.

### A possible role for plasticity

Despite evidence of genomic homogenisation in the savannah, morphological associations with environment were not weaker than in the fragmented rainforest. In fact, PEA models were consistently stronger, suggesting locally adaptive body shape variation. Although unexpected, this is not necessarily incongruous with homogenising gene flow, with phenotypic plasticity being a particularly favourable explanation. While G×P×E results indicate some heritable shape variation in both ecoregions, weaker associations in the savannah suggest partial decoupling of genotype and phenotype, consistent with plastic divergence (Schmid and Guillaume 2017). Supporting this, common garden studies in related rainbowfishes show that morphological plasticity can explain shape variation, including flow-induced divergence during development (McGuigan et al. 2003, Kelley et al. 2017).

Although regional differences in plasticity cannot be confirmed without experiments, alternative explanations (e.g., genotype–environment covariance; Conover and Schultz 1995) are possible. Still, theory predicts increased plasticity in systems with high gene flow and environmental heterogeneity (Sultan and Spencer 2002, Crispo 2008), suggesting savannah connectivity may favour plastic responses over local genetic specialisation. Conversely, in fragmented rainforest systems, plastic traits may become genetically assimilated (Pigliucci et al. 2006, Fitzpatrick 2012), potentially explaining stronger genotype–phenotype–environment associations. Plasticity may also be favoured under temporal variability, broadening tolerance to fluctuating environmental conditions (Janzen 1967, Hendry 2015). Overall, these results highlight testable hypotheses about the role of plasticity, not only for body shape but for broader physiological adaptation across rainforest and savannah systems.

## Conclusion

Tropical rainforest and savannah ecosystems in north-eastern Australia differ in key features: the former is ecologically stable and structurally complex, while the latter is more variable and highly connected. Accordingly, rainforest and savannah fish populations diverged not only in their responses to hydroclimatic variation, but also in the extent of local adaptation. Rainforest structure appears to promote diversification and specialisation, but may also require greater adaptive turnover under rapid change. In contrast, savannah populations show genomic homogenisation in a connected landscape, alongside notable phenotypic flexibility. These contrasting genomic and phenotypic patterns highlight the value of integrating multiple data types when studying selection. Beyond theory, such insights are important for effective management under ongoing environmental change.

## Acknowledgements

We acknowledge the Traditional Owners of Country covering the Wet Tropics, recognise their continuing connection to land, water and fishes, and pay respects to Elders past, present and emerging. We thank the members of MELFU and CEBEL at Flinders University and Labo Bernatchez at Laval University who helped with valuable theoretical discussions and methodological support. We thank Keith Martin for field support and discussions about rainbowfish ecology and distributions. Financial support was provided by the Australian Research Council (DP150102903 and FT130101068 to LBB), the Royal Society of South Australia for the Advancement of Science, and the Flinders University Student Association Development Grant. KG was supported by the AJ & IM Naylon PhD Scholarship via Flinders University, and the Playford Trust/Thyne Reid Foundation PhD Scholarship.

## Data Availability Statement

The relevant data have been appropriately archived and are available at figshare: https://figshare.com/s/c4b75d40fea91ab38beb. Data will be made publicly available upon acceptance.

## Ethical Approval

Animal handling procedures were performed in accordance with Flinders University Animal Ethical Approval E463/17. Field collection was performed under General fisheries Permit 191126 (Fisheries Act 1994; Queensland Government).

