## Supplemental Files for "Tropical rainforest *versus* savannah: Biome-specific landscape structure mediates adaptive strategies and resilience to environmental change"

Supplementary Material

Supplementary
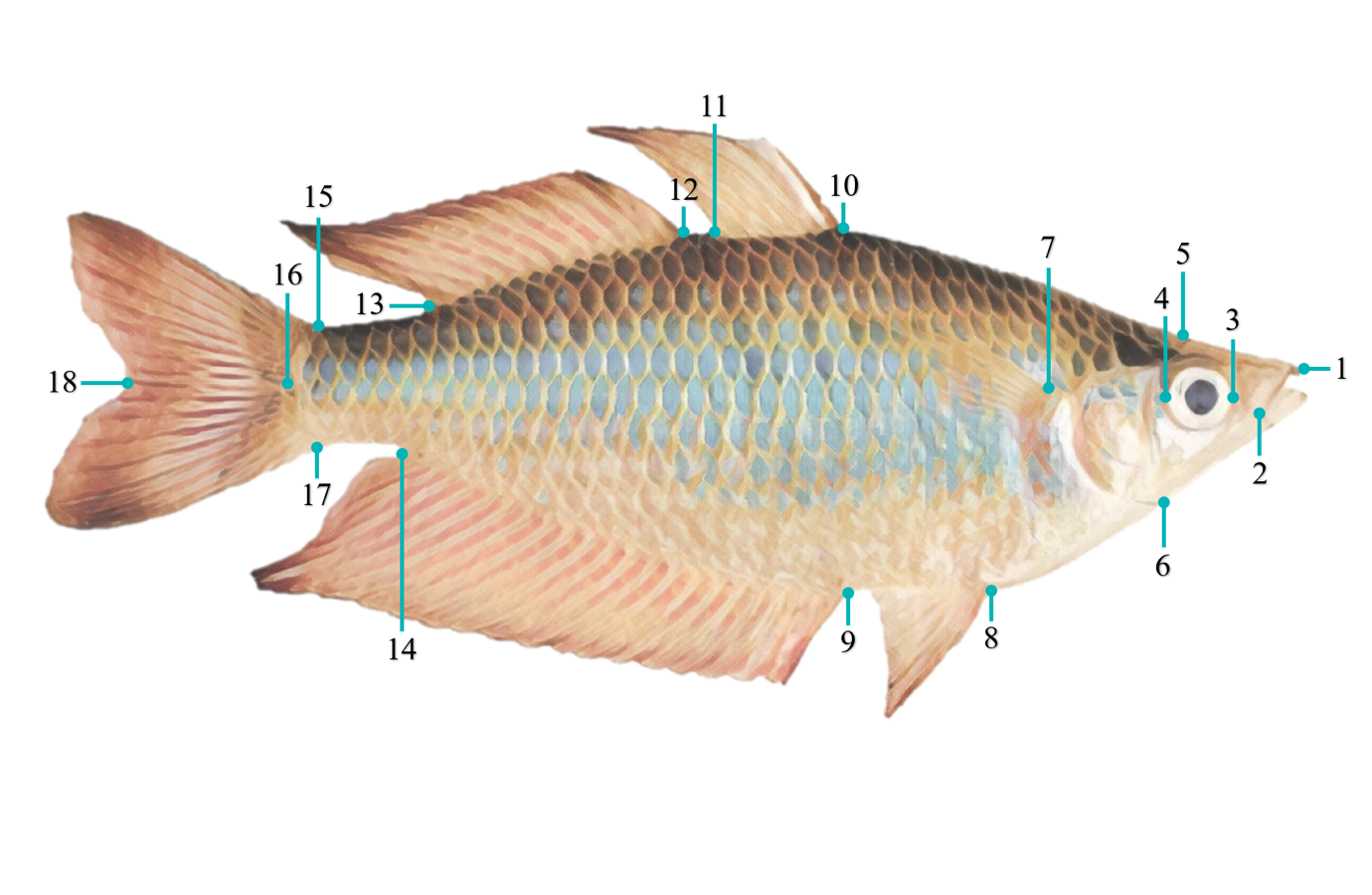
Figure 1. The 18 landmarks used for geometric morphometric analysis of the eastern rainbowfish *Melanotaenia splendida splendida*. 1: Anterior tip of head, where premaxillary bones articulate at midline; 2: Posterior tip of maxilla; 3: Anterior margin in maximum eye width; 4: Posterior margin in maximum eye width; 5: Dorsal margin of head at beginning of scales; 6: Ventral margin in the end of the head; 7: Dorsal insertion of pectoral fin; 8: Anterior insertion of the pelvic fin; 9: Anterior insertion of the anal fin; 10: Anterior insertion of the first dorsal fin; 11: Posterior insertion of the first dorsal fin; 12: Anterior insertion of the second dorsal fin; 13: Posterior insertion of the second dorsal fin; 14: Posterior insertion of the anal fin; 15: Dorsal insertion of the caudal fin; 16: Posterior margin of the caudal peduncle (at tip of lateral line); 17: Ventral insertion of the caudal fin; 18: Posterior margin of the caudal fin between dorsal and ventral lobes.

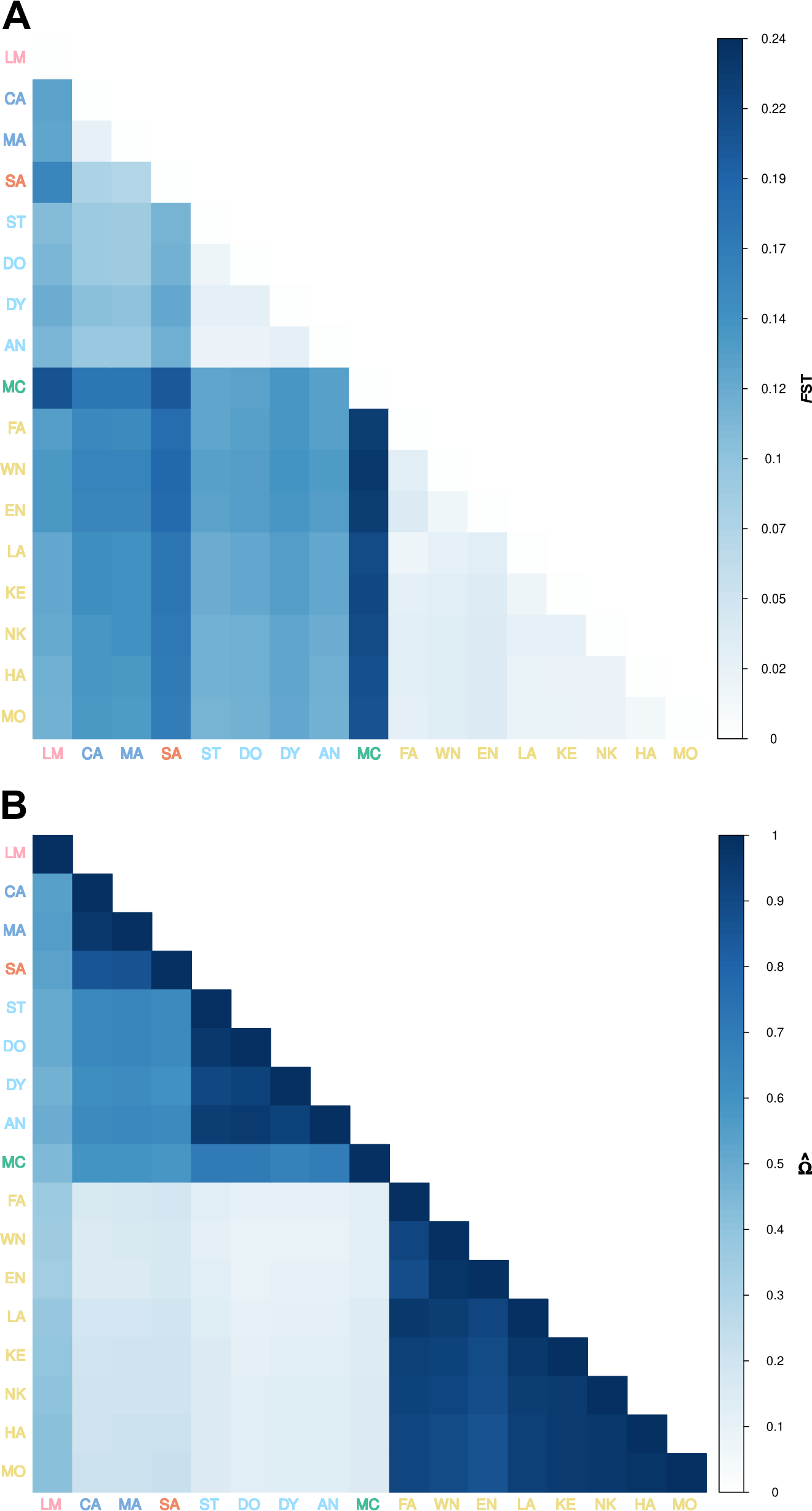

Supplementary Figure 2. Genomic differentiation and population structuring among rainforest and savannah sampling localities for the eastern rainbowfish *Melanotaenia splendida splendida*, represented by **(A)** Heatmap of pairwise F_ST_ based on 14,478 putatively neutral SNPs; **(B)** Correlation map for BAYPASS core model scaled covariance matrix Ω based on allele frequencies of the full dataset of 14,540 SNPs. Locality abbreviations follow Table 1, with colouration reflecting drainage of origin as in Figure 1.

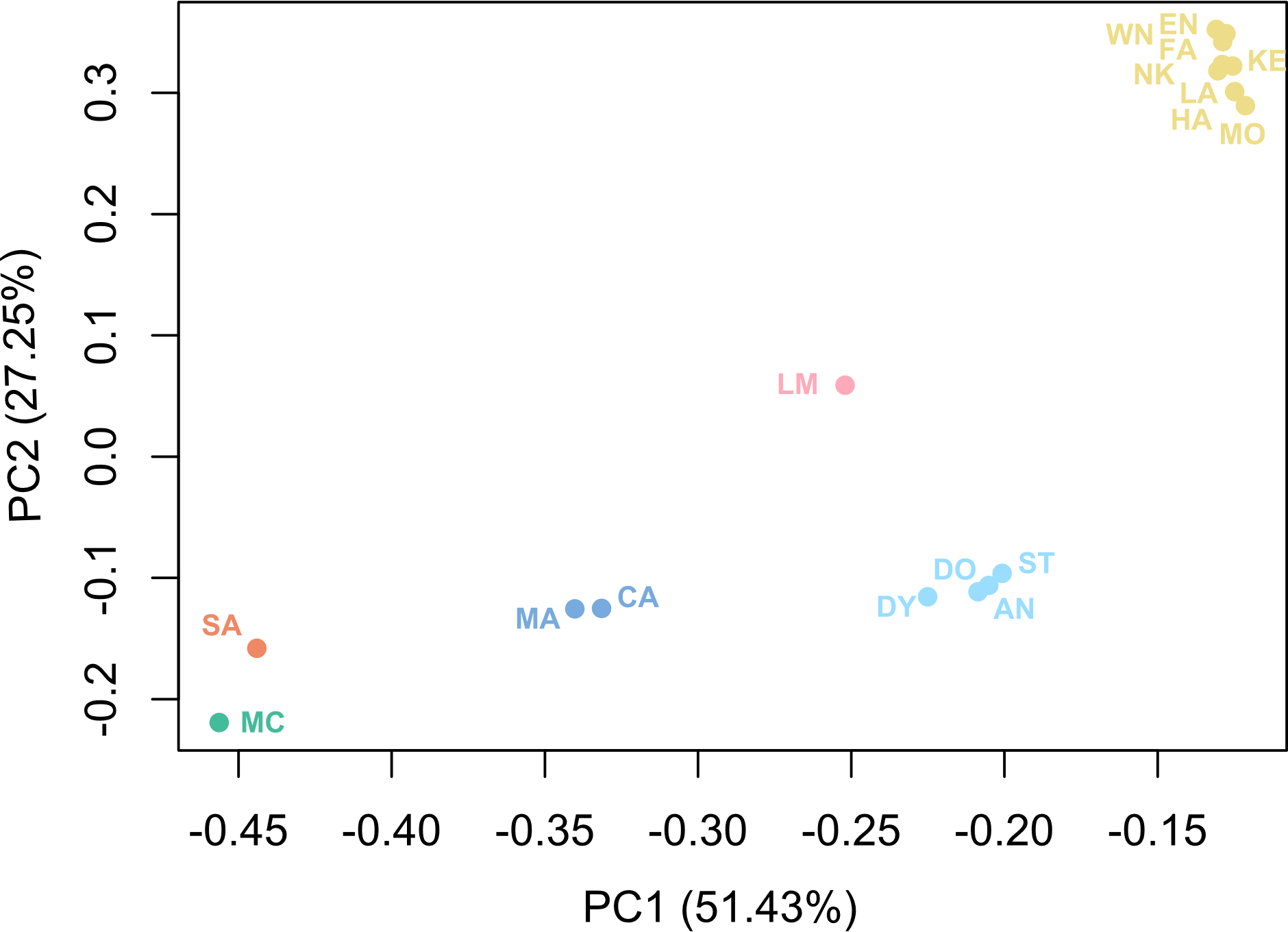

Supplementary Figure 3. Eigen-decomposition of scaled covariance matrix of locality-specific allele frequencies for *Melanotaenia splendida splendida*, based on 14,540 SNPs. Points correspond to sampling sites, and are colour coded by drainage system of origin following Figure 1, main text.

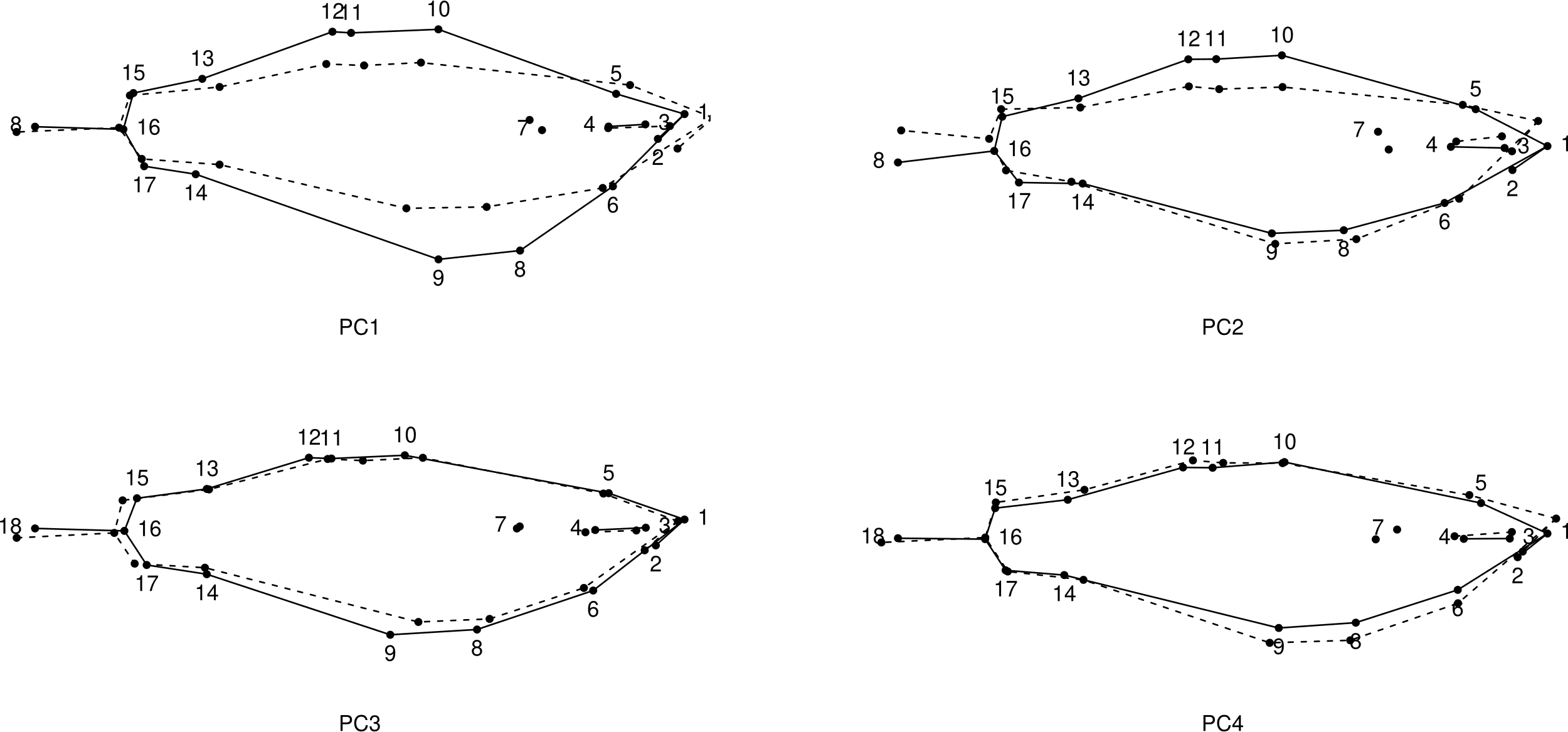

Supplementary Figure 4. Principal component analysis of body shape of all M. s. splendida individuals produced four significant PCs under broken stick modelling Wireframe graphical representation of significant principal components of body shape variation based on 18 landmarks for 366 *Melanotaenia splendida splendida* individuals sampled across seventeen rainforest and savannah sampling localities in tropical north-eastern Australia. Solid and dashed frames respectively represent body shape at high and low extremes of each significant axis (scale factor = 1).

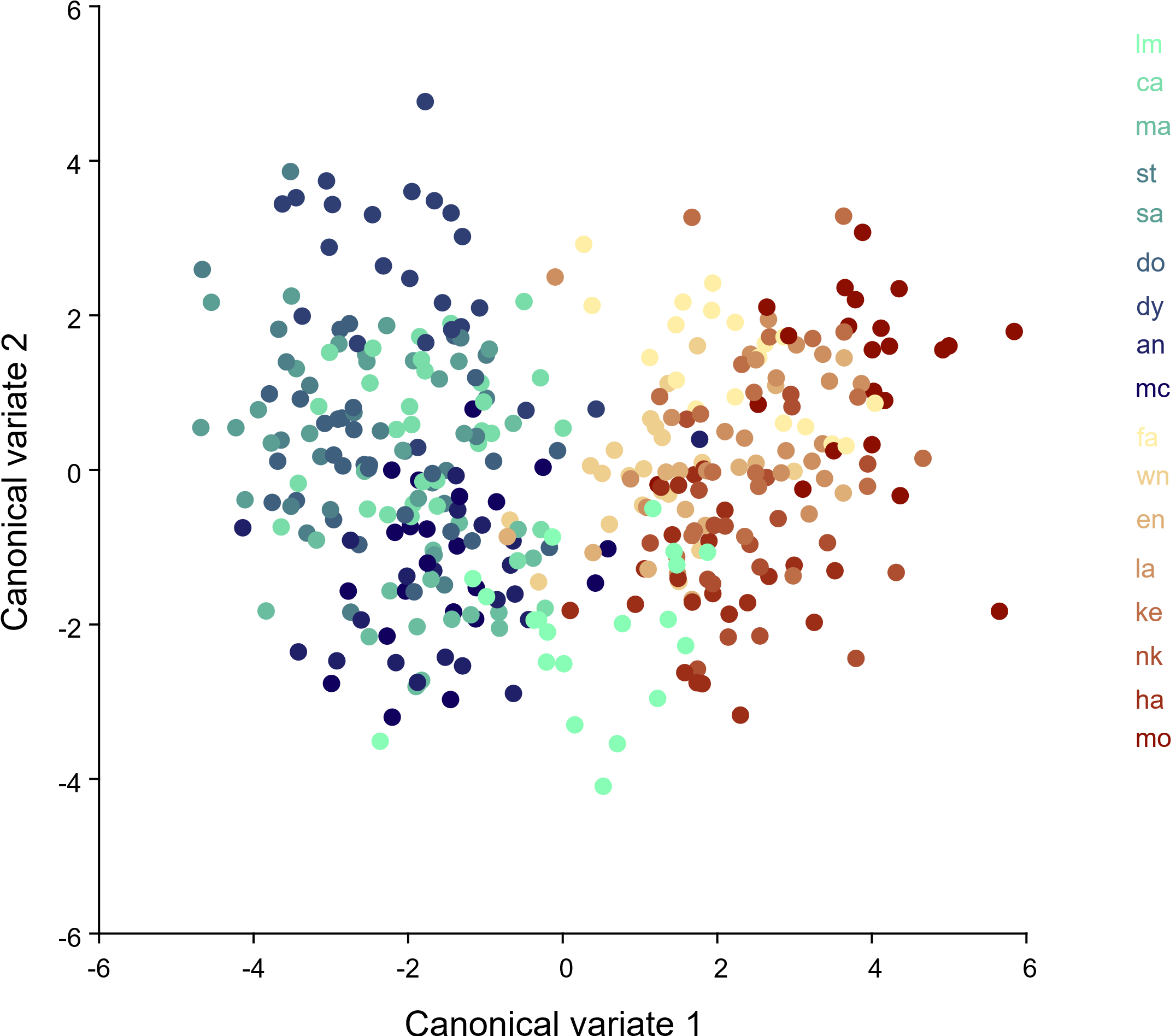

Supplementary Figure 5. Individual canonical body shape variation of 366 *Melanotaenia splendida splendida* individuals sampled across seventeen rainforest and savannah sampling localities in tropical north-eastern Australia. Locality codes follow Figure 1.

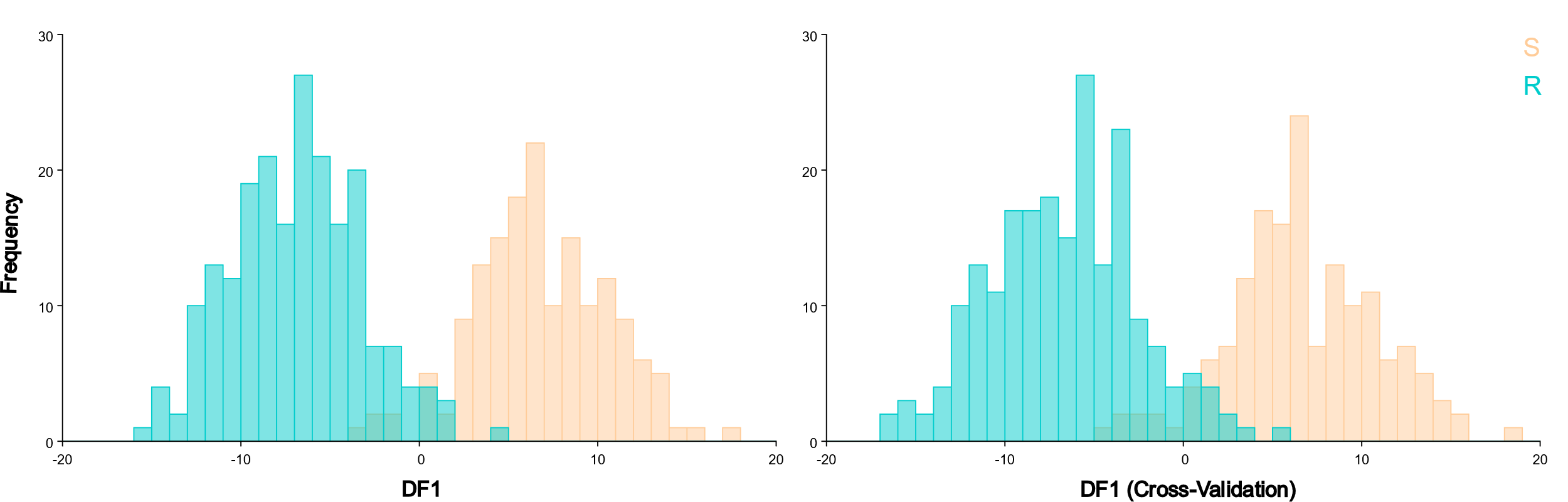

Supplementary Figure 6. Frequencies of individual discriminant function (DF) scores among rainforest (*n* = 207) and savannah (*n =*  159) *Melanotaenia splendida splendida*, based on multivariate analysis of 18 morphometric landmarks, controlling for centroid size.

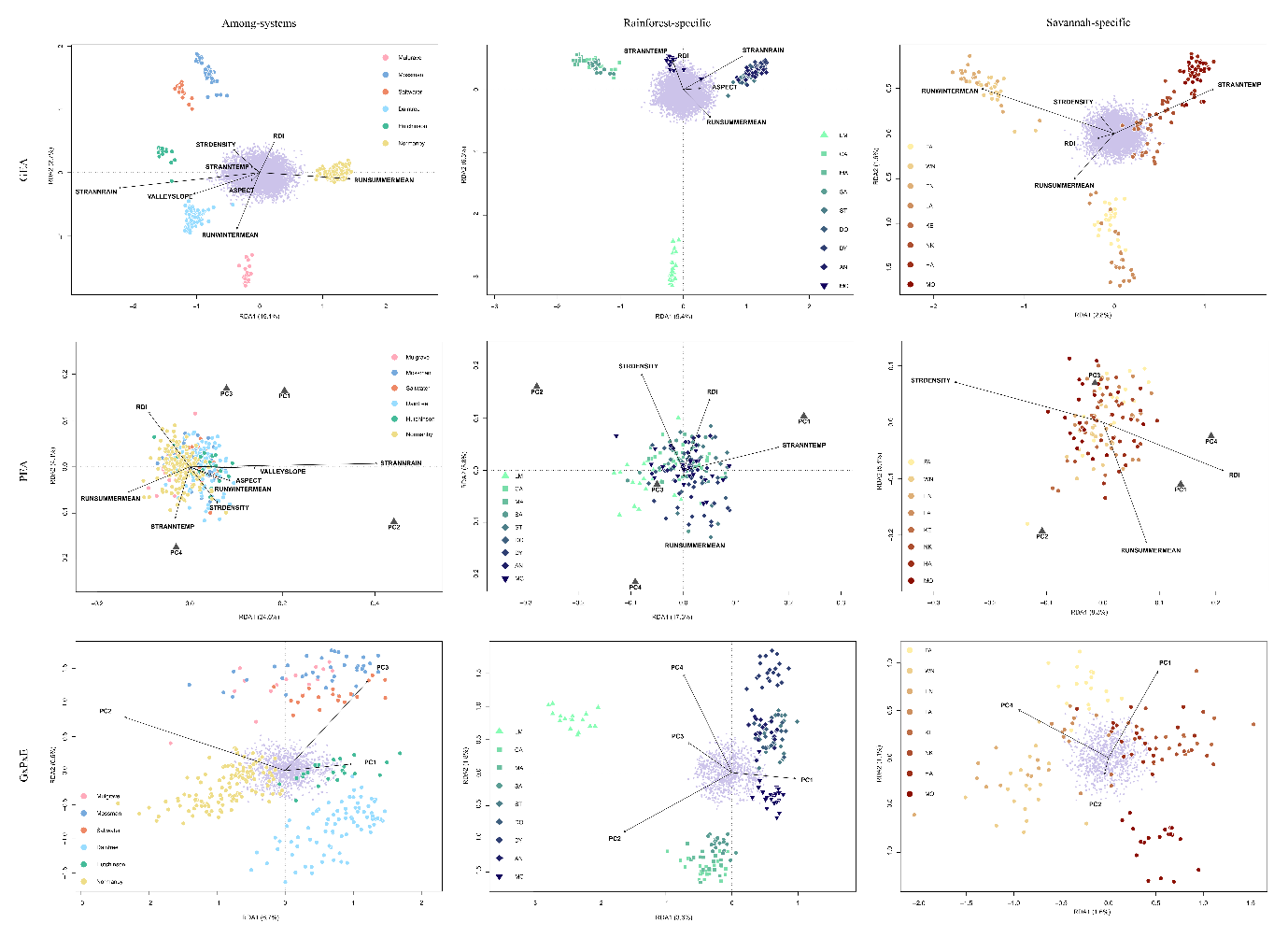

Supplementary Figure 7. Ordination plots summarising the first two axes of partial redundancy analyses (pRDAs) for *Melanotaenia splendida splendida* individuals sampled in tropical north-eastern Australia from 17 sampling localities (‘Between-systems’), including nine within the rainforest (‘rainforest-specific’) and eight within the savannah (‘savannah-specific’). Figures a-c represent genotype-environment associations (GEA) controlling for allelic covariance, d-f represent phenotype-environment associations (PEA) controlling for allelic covariance and body size, and g-i represent genotype-phenotype-environement associations (GxPxE) controlling for body size. Large points represent individual-level responses, and are coloured by drainage system of origin in ‘Between-systems’ plots, and by sampling site in ecoregion-specific plots. Small purple points represent SNP-level responses. Grey triangles represent morphological responses. Vectors represent the magnitude and direction of relationships with explanatory PCs.

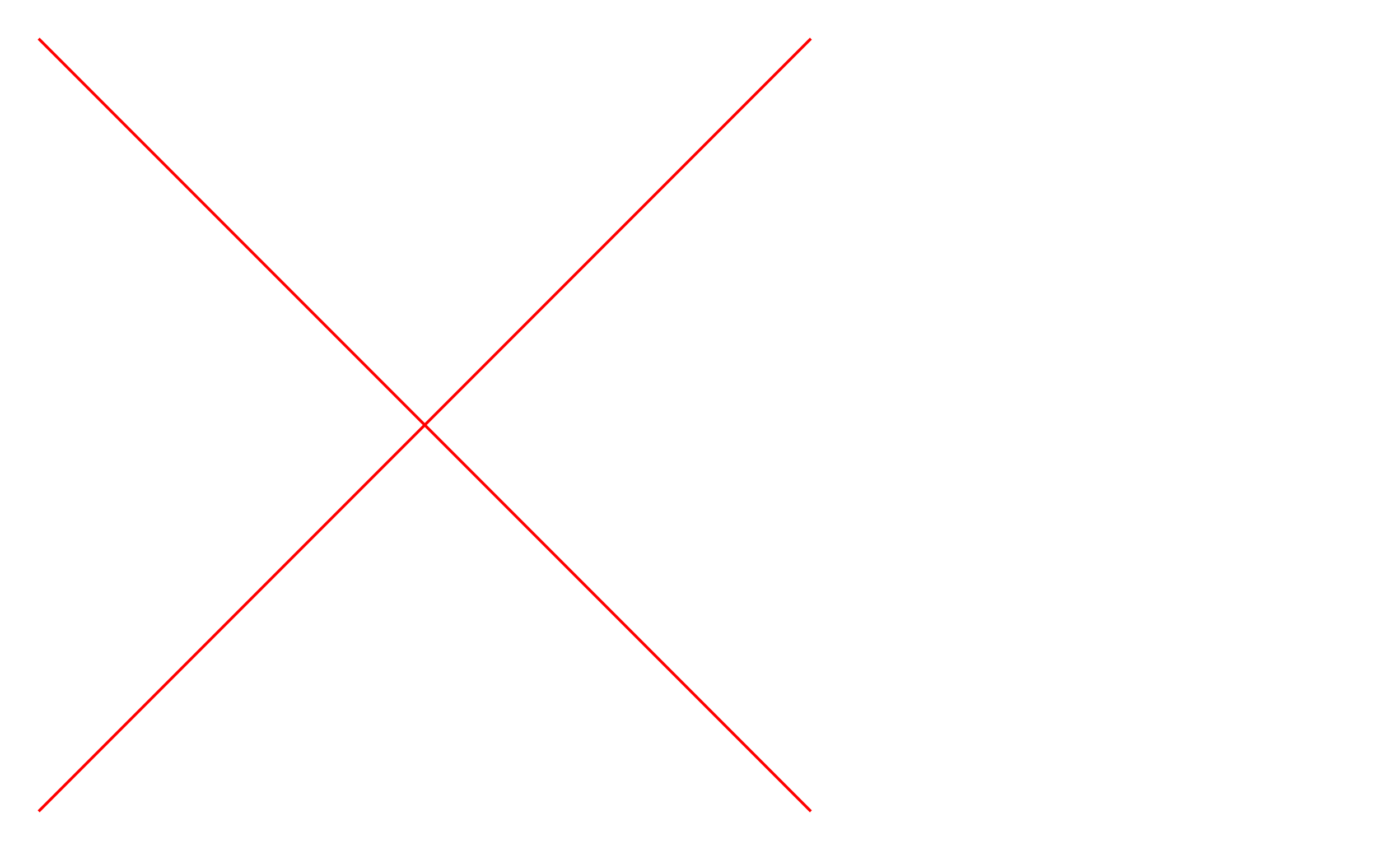

Supplementary Figure 8. Environmental association of 14,540 SNPs for *Melanotaenia splendida splendida* across seventeen sampling sites, against eight independent environmental variables, using BAYPASS auxiliary covariate model. Dashed line indicates Bayes Factor cutoff of 26 dB (99.8% probability), above which 233 loci were identified as candidates for environmental adaptation.

Supplementary Table 1. Raw climate data for each sampling locality of *Melanotaenia splendida splendida*. Shading represents relative variation among sites specific to each variable. Locality abbreviations: refer to Table 1 (main text). Compiled from Stein, J. L., Hutchison, M.F., Stein, J.A. 2011. National Environmental Stream Attributes v1.1.3. Page <http://pid.geoscience.gov.au/dataset/ga/73045> Geoscience Australia, Canberra. Accessed June 2017.

|  | STR ANN TEMP (**°**C) | STR ANN RAIN (MM) | RUN SUMMER MEAN (mL) | RUN WINTER MEAN (mL) | RDI (index: 0-1) | VALLEY SLOPE | ASPECT (**°**) | STR DENSITY (km/km^2^) |
| --- | --- | --- | --- | --- | --- | --- | --- | --- |
| LM | 23.95 | 2025.63 | 32734.99 | 841.00 | 0.01 | 0.85 | 127.01 | 1.01 |
| SA | 24.47 | 203.50 | 8630.67 | 14.67 | 0.07 | 0.32 | 338.10 | 0.97 |
| MA | 24.46 | 221.71 | 5887.23 | 10.44 | 0.12 | 0.62 | 25.30 | 0.93 |
| SA | 24.33 | 2350.00 | 11915.01 | 153.05 | 0.02 | 0.22 | 51.90 | 1.14 |
| ST | 24.40 | 2742.62 | 51975.84 | 601.22 | 0.01 | 0.12 | 340.19 | 1.03 |
| DO | 24.33 | 2764.57 | 31506.39 | 205.31 | 0.02 | 0.43 | 60.76 | 1.08 |
| DY | 24.33 | 3173.78 | 4013.73 | 44.02 | 0.05 | 5.51 | 142.07 | 0.57 |
| AN | 24.13 | 3153.86 | 2036.09 | 21.18 | 0.15 | 9.24 | 212.48 | 0.79 |
| MC | 23.73 | 3181.89 | 1766.47 | 21.67 | 0.05 | 9.41 | 15.07 | 1.22 |
| FA | 21.77 | 1129.53 | 2166.42 | 0.33 | 0.06 | 4.04 | 18.59 | 0.81 |
| WN | 23.10 | 1308.29 | 305467.60 | 345.83 | 0.05 | 0.20 | 354.11 | 1.01 |
| EN | 23.08 | 1381.96 | 130284.80 | 241.02 | 0.09 | 0.13 | 337.75 | 0.94 |
| LA | 24.70 | 976.38 | 45350.50 | 12.93 | 0.08 | 0.06 | 17.35 | 0.95 |
| KE | 25.20 | 1017.09 | 259012.90 | 0.00 | 0.04 | 0.04 | 14.31 | 0.90 |
| NK | 25.31 | 1044.46 | 64037.12 | 0.00 | 0.03 | 0.07 | 27.78 | 0.63 |
| HA | 25.30 | 107.00 | 173939.90 | 0.00 | 0.10 | 0.18 | 62.61 | 0.61 |
| MO | 25.40 | 1073.47 | 341.29 | 0.00 | 0.05 | 0.37 | 109.16 | 1.38 |

Supplementary Table 2. Pairwise *F*_ST_ among *Melanotaenia splendida splendida* from 17 sampling sites across in tropical north-eastern Australia, based on 14,478 putatively neutral SNPs. Locality abbreviations: refer to Table 1, main text.

|  | CA | MA | SA | ST | DO | DY | AN | MC | FA | WN | EN | LA | KE | NK | HA | MO |
| --- | --- | --- | --- | --- | --- | --- | --- | --- | --- | --- | --- | --- | --- | --- | --- | --- |
| CA | 0.128 |  |  |  |  |  |  |  |  |  |  |  |  |  |  |  |
| MA | 0.126 | 0.026 |  |  |  |  |  |  |  |  |  |  |  |  |  |  |
| SA | 0.158 | 0.075 | 0.071 |  |  |  |  |  |  |  |  |  |  |  |  |  |
| ST | 0.105 | 0.087 | 0.084 | 0.111 |  |  |  |  |  |  |  |  |  |  |  |  |
| DO | 0.108 | 0.088 | 0.086 | 0.113 | 0.017 |  |  |  |  |  |  |  |  |  |  |  |
| DY | 0.119 | 0.099 | 0.097 | 0.125 | 0.029 | 0.027 |  |  |  |  |  |  |  |  |  |  |
| AN | 0.109 | 0.090 | 0.089 | 0.115 | 0.021 | 0.019 | 0.027 |  |  |  |  |  |  |  |  |  |
| MC | 0.208 | 0.174 | 0.174 | 0.202 | 0.127 | 0.130 | 0.141 | 0.132 |  |  |  |  |  |  |  |  |
| FA | 0.134 | 0.155 | 0.153 | 0.184 | 0.127 | 0.131 | 0.141 | 0.131 | 0.227 |  |  |  |  |  |  |  |
| WN | 0.138 | 0.159 | 0.159 | 0.188 | 0.129 | 0.134 | 0.144 | 0.135 | 0.232 | 0.030 |  |  |  |  |  |  |
| EN | 0.144 | 0.165 | 0.166 | 0.194 | 0.134 | 0.138 | 0.148 | 0.139 | 0.237 | 0.040 | 0.019 |  |  |  |  |  |
| LA | 0.124 | 0.146 | 0.145 | 0.174 | 0.118 | 0.123 | 0.132 | 0.123 | 0.215 | 0.017 | 0.024 | 0.032 |  |  |  |  |
| KE | 0.124 | 0.147 | 0.146 | 0.175 | 0.118 | 0.123 | 0.133 | 0.124 | 0.218 | 0.027 | 0.029 | 0.037 | 0.019 |  |  |  |
| NK | 0.121 | 0.142 | 0.145 | 0.172 | 0.113 | 0.117 | 0.127 | 0.119 | 0.214 | 0.029 | 0.033 | 0.041 | 0.025 | 0.024 |  |  |
| HA | 0.116 | 0.139 | 0.138 | 0.168 | 0.113 | 0.117 | 0.127 | 0.117 | 0.210 | 0.029 | 0.033 | 0.041 | 0.023 | 0.021 | 0.020 |  |
| MO | 0.115 | 0.138 | 0.136 | 0.166 | 0.111 | 0.114 | 0.124 | 0.115 | 0.209 | 0.029 | 0.033 | 0.041 | 0.024 | 0.023 | 0.020 | 0.014 |

Supplementary Table 3. Procrustes distances among sampling sites, based on canonical variate analysis of body shape of *Melanotaenia splendida splendida* across rainforest and savannah localities (For codes refer to Figure 1). Significant differences (*p* = <0.05) are indicated by bold font.

|  | AN | CA | DO | DY | EN | FA | HA | KE | LA | LM | MA | MC | MO | NK | SA | ST |
| --- | --- | --- | --- | --- | --- | --- | --- | --- | --- | --- | --- | --- | --- | --- | --- | --- |
| CA | 0.0138 |  |  |  |  |  |  |  |  |  |  |  |  |  |  |  |
| DO | 0.0142 | 0.0165 |  |  |  |  |  |  |  |  |  |  |  |  |  |  |
| DY | 0.0256 | 0.0266 | 0.0204 |  |  |  |  |  |  |  |  |  |  |  |  |  |
| EN | 0.0282 | 0.0234 | 0.0339 | 0.0403 |  |  |  |  |  |  |  |  |  |  |  |  |
| FA | 0.0338 | 0.0277 | 0.0391 | 0.0441 | 0.0139 |  |  |  |  |  |  |  |  |  |  |  |
| HA | 0.0288 | 0.0243 | 0.0346 | 0.0412 | 0.012 | 0.0147 |  |  |  |  |  |  |  |  |  |  |
| KE | 0.0321 | 0.0275 | 0.0355 | 0.0337 | 0.0186 | 0.0202 | 0.0193 |  |  |  |  |  |  |  |  |  |
| LA | 0.0263 | 0.0214 | 0.0304 | 0.0322 | 0.0151 | 0.0197 | 0.0152 | 0.0101 |  |  |  |  |  |  |  |  |
| LM | 0.0312 | 0.0289 | 0.0351 | 0.0368 | 0.0203 | 0.0267 | 0.0224 | 0.0186 | 0.0199 |  |  |  |  |  |  |  |
| MA | 0.0138 | 0.0148 | 0.0156 | 0.0275 | 0.0247 | 0.0324 | 0.0254 | 0.0288 | 0.0234 | 0.0243 |  |  |  |  |  |  |
| MC | 0.008 | 0.014 | 0.012 | 0.0241 | 0.0294 | 0.035 | 0.0297 | 0.032 | 0.026 | 0.0313 | 0.0134 |  |  |  |  |  |
| MO | 0.0396 | 0.0354 | 0.0446 | 0.0412 | 0.0244 | 0.0229 | 0.0248 | 0.0121 | 0.0192 | 0.0263 | 0.0383 | 0.0403 |  |  |  |  |
| NK | 0.0372 | 0.0324 | 0.0415 | 0.0442 | 0.0159 | 0.0163 | 0.0176 | 0.0152 | 0.0185 | 0.0197 | 0.032 | 0.0379 | 0.0183 |  |  |  |
| SA | 0.0235 | 0.0237 | 0.0151 | 0.0153 | 0.0406 | 0.0445 | 0.0408 | 0.0362 | 0.0334 | 0.0384 | 0.0238 | 0.0209 | 0.0446 | 0.0451 |  |  |
| ST | 0.0184 | 0.0244 | 0.0129 | 0.0222 | 0.0424 | 0.0476 | 0.0436 | 0.0428 | 0.0378 | 0.0434 | 0.0243 | 0.0163 | 0.0508 | 0.0501 | 0.0161 |  |
| WN | 0.0176 | 0.0175 | 0.0224 | 0.0247 | 0.0194 | 0.0244 | 0.0217 | 0.0178 | 0.0152 | 0.0221 | 0.0187 | 0.0184 | 0.0249 | 0.0254 | 0.026 | 0.0285 |

Supplementary Table 4. Classification/misclassification tables for discriminant function analysis of 18 morphometric landmarks among rainforest (*n* = 207) and savannah (*n =* 159) *Melanotaenia splendida splendida*, controlling for centroid size. Differences between means = 0.0274 Procrustes distance, 3.705 Mahalanobis distance. *P*-value (parametric) = <.0001 for 1000 permutation runs.

| Discriminant function | True origin | Allocated to | | |
| --- | --- | --- | --- | --- |
|  |  | Rainforest | Savannah | Total |
|  | Rainforest | 200 | 8 | 208 |
|  | Savannah | 5 | 154 | 159 |
| Cross-validation | True origin | Allocated to | | |
|  |  | Rainforest | Savannah | Total |
|  | Rainforest | 195 | 13 | 208 |
|  | Savannah | 8 | 151 | 159 |

Supplementary Table 5. Model significance for pRDAs testing genotype-environment (GEA), phentoype-environment (PEA), and genotyoe-phenotype-environment (GxPxE) associations for *Melanotaenia spplendida splendida*, based on ANOVA-like permutation test for Constrained Correspondence Analysis (anova.cca) using 999 permutation rounds.

| Analysis | Response Dataset | Covariable | Df | | Variance | | F |
| --- | --- | --- | --- | --- | --- | --- | --- |
|  |  |  | Model | Residual | Model | Residual |  |
| GEA | Among-ecotypes | Allelic covariance | 8 | 369 | 889.75 | 3076.13 | 13.341*** |
|  |  | *F*_ST_ | 8 | 370 | 357.33 | 3042.78 | 5.431*** |
|  |  | river dist | 8 | 370 | 376.59 | 3051.43 | 5.708*** |
|  | Rainforest-specific | Allelic covariance | 5 | 201 | 625.55 | 2745.1 | 9.161*** |
|  |  | *F*_ST_ | 6 | 201 | 456.47 | 2745.1 | 5.571*** |
|  | Savannah-specific | Allelic covariance | 5 | 163 | 223.88 | 3041.22 | 2.3999*** |
|  |  | *F*_ST_ | 5 | 163 | 126.18 | 3041.22 | 1.3525*** |
| PEA | Among-ecotypes | Allelic covariance + size | 6 | 294 | 0.000187 | 0.000397 | 23.03*** |
|  |  | *F*_ST_ + size | 7 | 294 | 0.000086 | 0.000402 | 8.939*** |
|  |  | River distance + size | 7 | 294 | 0.000086 | 0.000402 | 8.939*** |
|  | Rainforest-specific | Allelic covariance + size | 4 | 171 | 0.000159 | 0.000486 | 13.969*** |
|  |  | *F*_ST_ + size | 4 | 172 | 0.000137 | 0.000494 | 11.972*** |
|  | Savannah-specific | Allelic covariance + size | 3 | 117 | 0.000062 | 0.000385 | 6.2968*** |
|  |  | *F*_ST_ + size | 4 | 117 | 0.000077 | 0.000382 | 5.8896*** |
| GxPxE | Among-ecotypes | Size | 3 | 297 | 44.01 | 527.96 | 8.253*** |
|  | Rainforest-specific | Size | 4 | 171 | 28.27 | 404.62 | 2.9866*** |
|  | Savannah-specific | Size | 3 | 120 | 14.45 | 396.92 | 1.4563*** |

Supplementary Table 6. Proportion of variation better explained by environment versus neutral factors in patrial RDAs

|  | Controlling for | Genomic | Morphological |
| --- | --- | --- | --- |
| Among-ecotypes | Allelic covariance | 5 | 12 |
|  | *F*_ST_ | 4 | 4 |
|  | River distance | 2 | 9 |
| Rainforest-specific | Allelic covariance | 4 | 19 |
|  | *F*_ST_ | 3 | 19 |
| Savannah-specific | Allelic covariance | 3 | 5 |
|  | *F*_ST_ | 2 | 6 |
|  | Average | 3.3 | 10.5 |
|  | Standard Deviation | 1.0 | 5.9 |
